# Hierarchical structural organization in bioinspired peptide coacervate microdroplets

**DOI:** 10.1101/2024.07.06.602323

**Authors:** Jessica Lim, Sushanth Gudlur, Claire Buchanan, Quentin Moana Perrin, Hannah Boyd, Martine Moulin, Hiroki Iwase, Lionel Porcar, Marité Cárdenas, Ali Miserez, Konstantin Pervushin

## Abstract

This study explores the dynamic and hierarchical structural organization of peptide coacervate microdroplets at the meso-to atomic-scale resolution using a combination of Transferred Nuclear Overhauser Effect Spectroscopy (TrNOESY), Small Angle Neutron Scattering (SANS), and confocal microscopy. Dynamic interactions driving the self-association of peptide clusters are revealed, highlighting the critical roles of interacting residues. These phase-separating model peptides form small oligomers at low pH, which aggregate into larger clusters at neutral pH. These clusters organize into a porous network within the droplets, facilitating size-selective cargo sequestration. The findings underscore the significance of the dynamic spatio-temporal properties of peptide-based coacervates, contributing to our understanding of phase separation at the atomic and molecular levels. Critically, this approach enables the investigation of coacervate structures in their native state, offering insights into the physical and dynamic interactions governing droplet formation and cargo encapsulation.

**TOC:** 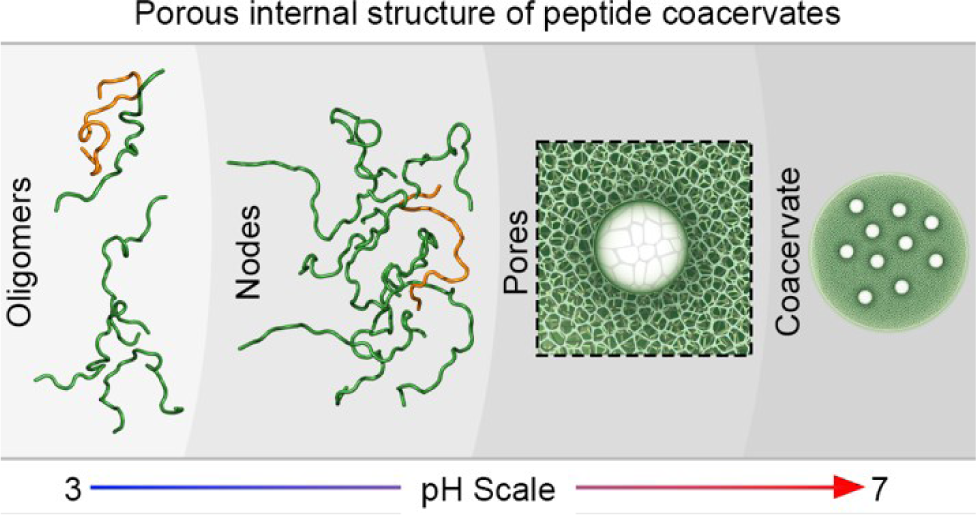

## INTRODUCTION

Investigating the internal structure of coacervates (biocondensates) assembled by liquid-liquid phase separation (LLPS) is challenging due to their complex and dynamic nature. Imaging techniques such as scanning electron microscopy (SEM), transmission electron microscopy (TEM), and cryo-electron tomography have enabled the visualization of their diverse internal architectures ranging from amorphous assemblies to structured complexes and lateral biopolymer interactions, but these techniques fail to capture the dynamic transient states within the droplet microenvironment^1–4^. Likewise, most optical microscopy techniques are limited in their spatiotemporal resolution^5^. While super-resolution microscopy techniques may significantly improve spatial resolution and molecule localization microscopy (SMLM) was recently employed to reveal spatial inhomogeneities within the coacervate interior^6^, one is limited by the available inventory of fluorogens. Computational modelling has provided valuable insights into the structural organization within coacervates^7–9^, but is limited by the need to simulate extremely large ensembles of the self-organizing building blocks, where even slight adjustments in the simulation parameters can significantly alter the resulting structures. To overcome the above mentioned limitations, more versatile techniques are needed to capture the full range of structural dynamics within the coacervate interior.

NMR spectroscopy is a powerful tool for studying peptide structures, even when they are highly mobile and dynamic^10^. NMR spectroscopy excels in capturing the conformational flexibility and transient interactions characteristic of such dynamic systems^10,11^. It provides detailed atomic-level information on the peptide backbone and side-chain conformations, enabling the elucidation of structural ensembles that reflect the full range of motion and spatial heterogeneity. This capability is particularly valuable for investigating intrinsically disordered peptides and proteins, which do not adopt a single static structure but instead exist as an ensemble of rapidly interconverting conformations^10–12^. However, NMR in both solution and solid-state faces significant limitations when applied to LLPS due to their high viscosity or structural inhomogeneity, respectively. The high viscosity of coacervate systems results in slow molecular tumbling and reduced rotational diffusion, which leads to broader NMR spectral lines and decreased resolution. This broadening of spectral lines impedes the ability to obtain clear, high-resolution spectra necessary for detailed structural analysis. Additionally, the extremely increased viscosity by several orders of magnitude in comparison to aqueous peptide solutions^13^ can result in longer relaxation times, complicating the acquisition of multidimensional NMR data and reducing the sensitivity and accuracy of the measurements. To address these limitations, here we employed a specialized NMR technique, Transferred Nuclear Overhauser Effect Spectroscopy (TrNOESY)^14,15^ in combination with confocal fluorescence microscopy and Small Angle Neutron Scattering (SANS) coupled with selective deuteration to investigate the internal structure of coacervates.

TrNOESY is a specialized NMR technique employed to investigate interactions and structures of ligands forming complexes with larger macromolecules, such as proteins or nucleic acids. It is particularly useful when the ligand rapidly exchanges between the free and bound states. In our TrNOESY experiments, the ligand of interest is a peptide capable of phase separating into coacervate microdroplets and exists in equilibrium between its free (dilute phase) and bound (dense phase or coacervated) states. When a free peptide interacts with any bound peptide in the dense phase, the NOE signals are transferred from the bound state to the free state. This transferred NOE (TrNOE) helps detect interactions even at low concentrations of the free peptides or when the complex is in rapid exchange. TrNOE is observed due to the significant difference in the rotational correlation times of the peptide between the free and bound states, where the slower tumbling in the latter enhances the NOE signals.

By leveraging the intrinsic material exchange between the dilute and dense phases of coacervates, TrNOESY experiments reveal dynamic interactions as the system approaches phase separation. These insights highlight residue-level interactions driving the self-association of peptide clusters and the critical roles of sticker and spacer residues. Our TrNOESY experiments are also consistent with the presence of nano-porous structures within the droplets, supporting the concept of spatial inhomogeneity in the microdroplet interior. Additionally, we propose that TrNOEs are detectable only if coacervates exhibit an extended network of internal pores or cavities facilitating peptide exchange between phases. This network was indirectly confirmed through confocal microscopy, which showed that the microdroplets act as molecular sieves, selectively allowing cargo of specific sizes into the microdroplet interior. Additionally, the granularity and dimensions of the internal structures were directly estimated by selective deuteration and contrast matched SANS.

## RESULTS

### Model peptides employed in this study and their phase behaviour

To study the internal structure of coacervates, we employed peptides from the HB*pep* family of peptides. The peptide sequences from this family are inspired by the histidine-rich beak proteins of the Humboldt squid and feature varying repeats of the GHGxY-type (where x is a hydrophobic residue) or GAGFA-type motifs^13,16^. Under physiological conditions, these peptides readily phase separate into ‘simple coacervates’ (composed of a single type of peptide). For this study, we selected three peptide variants (Fig. 1a) with differing phase separation propensities: weak (GY23) and strong ((GY23)_2_ & GW26), to capture any structural differences arising from their slight differences in peptide sequences or overall length.

**Figure 1.**
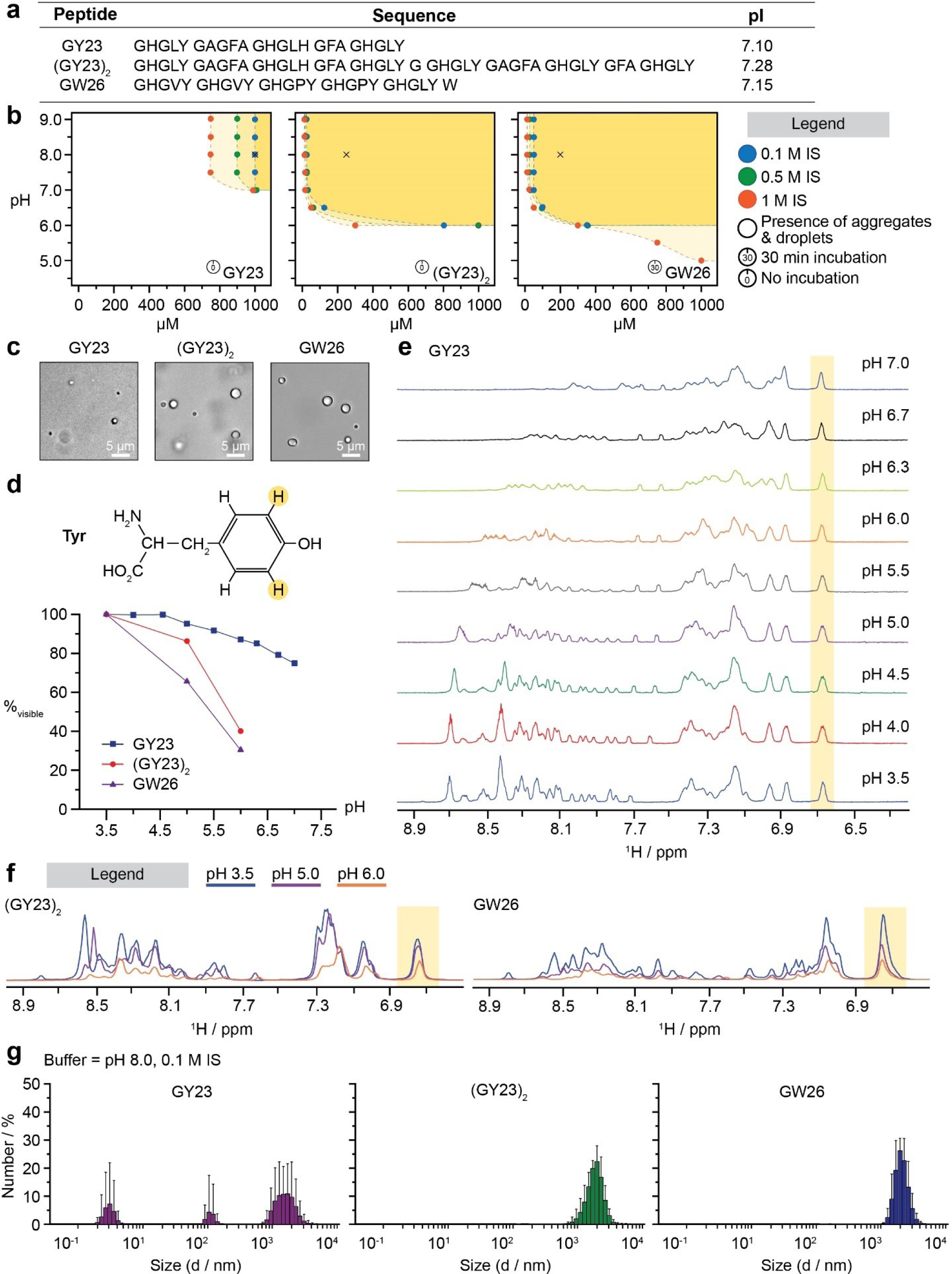
Phase behaviour of HB*pep* variants GY23, (GY23)_2_ and GW26. **(a)** Peptide sequences used in this study and their theoretical isoelectric point (IEP) calculated using ExPASY ProtParam^17^. **(b)** Phase diagrams of pH *vs.* peptide concentration for each peptide. Colours represent different ionic strengths (IS) as detailed in the legend. Points with black outlines indicate both aggregates and droplets visible under the microscope. Incubation time, if applicable, is shown in the legend. **(c)** Brightfield micrographs of peptides at pH 8.0, 0.1 M IS, corresponding to the black crosses in (b). **(d)** Chemical structure of Tyr where coloured H atoms correspond to highlighted peak in (e) and (f). Integrals from the identified (highlighted) peak across different pH conditions were normalised to 100 % at pH 3.5 (starting pH). All three peptides are plotted on the same graph for comparison. Lines are included for visual aid. **(e)** Broadening of proton (^1^H) resonances as GY23 approached phase separation at neutral pH. Measurements and corresponding data were obtained in a previous publication^18^. **(f)** Similar broadening of resonances for both (GY23)_2_ and GW26 was observed over indicated pH values. Overlaid spectra with baseline alignment to facilitate comparison. Only part of the spectra – amide and aromatic protons is depicted for both (e) and (f). **(g)** Number-based size distribution obtained from DLS measurements. Corresponding intensity- and volume-based plots can be found in Fig. S1. Concentrations used correspond to that for (c). Measurement details and data analysis can be found in Methods.

The three peptides were similar in their ability to spontaneously phase separate at or near neutral pH and in physiologically relevant ionic strength (IS) conditions (Fig. 1b). However, they differed slightly in their phase separation propensities, especially when the pH and ionic strength was varied. This was confirmed using brightfield microscopy (Fig. 1b, c) and dynamic light scattering (DLS) measurements (Fig. 1g). GY23 coacervated into relatively smaller droplets and within a narrow pH and IS regime while (GY23)_2_ – comprising two GY23 sequences linked by a glycine residue – and GW26 formed relatively large droplets under a wider pH and IS regime (Fig. 1c, g).

### 1H resonances of peptides in the dense phase are not detectable by NMR in solution

A series of one-dimensional (1D) experiments were conducted on the three peptides in the solution state whose individual concentrations were carefully chosen based on their phase diagrams (Fig. 1b) to avoid turbid samples caused by coacervation, thus ensuring good spectral quality. As expected, as the pH approached neutral values, the chemical shifts of the amide proton resonances progressively decreased in range and amplitude (Fig. 1e, f) due to the loss of secondary structure in the free peptides and accelerated exchange with bulk water, respectively^19,20^. However, the side chain protons of aromatic tyrosine (Tyr), which do not exchange with bulk water and are not attenuated by this exchange, still showed a signal decrease without significant resonance line broadening (Fig. 1d). This decrease is attributed to peptide sequestration into the dense phase. The tracked Tyr cross-peak integrals showed the most significant decrease in GW26, which had the highest phase separating propensity, followed by (GY23)_2_ and GY23 under similar experimental conditions, in agreement with their phase diagrams (Fig. 1b). Therefore, this signal could be used to quantitatively track the distribution of peptides in the NMR-observable phase of free peptides in solution and the NMR “dark” phase of peptides in the droplets.

### Detectability of TrNOE signals in coacervating peptides in equilibrium with the dense phase

TrNOESY is commonly employed to assess structural information of smaller molecular weight (MW) ligands bound to larger receptors or lipid vesicles that are typically inaccessible to solution-state NMR^14,15^. This method consists of the transfer of modulated spin magnetization from the ligand’s bound state to its free state, enabling the acquisition of structural data of the ligand in the bound state. Several conditions need to be met in order to detect TrNOEs in exchanging systems, the most important of which are the fast exchange reaction kinetic rates, high reaction equilibrium constant, reasonable equilibrium concentration of ligand, sufficiently available binding sites and sufficiently different rotational correlation times of the free and bound ligands^15,21,22^. To detect TrNOEs, manifested as the significant increase of the NOE cross-peak intensities relative to the diagonal, we systematically acquired NOESY spectra at different pH for the three peptides. As shown in Fig. 2a, we detected a more than tenfold increase in the relative NOE intensities when the pH approached phase separation conditions. TrNOEs were confirmed in a series of 2D NOESY spectra measured at various mixing times, exhibiting the build-up of TrNOEs at different pH conditions (Fig. 2b & Fig. S2). As no saturation of NOE intensity was observed across the mixing times explored, all NOEs were fitted with a linear regression, and the derived slope values were plotted across the peptide sequence in Fig. S3.

**Figure 2.**
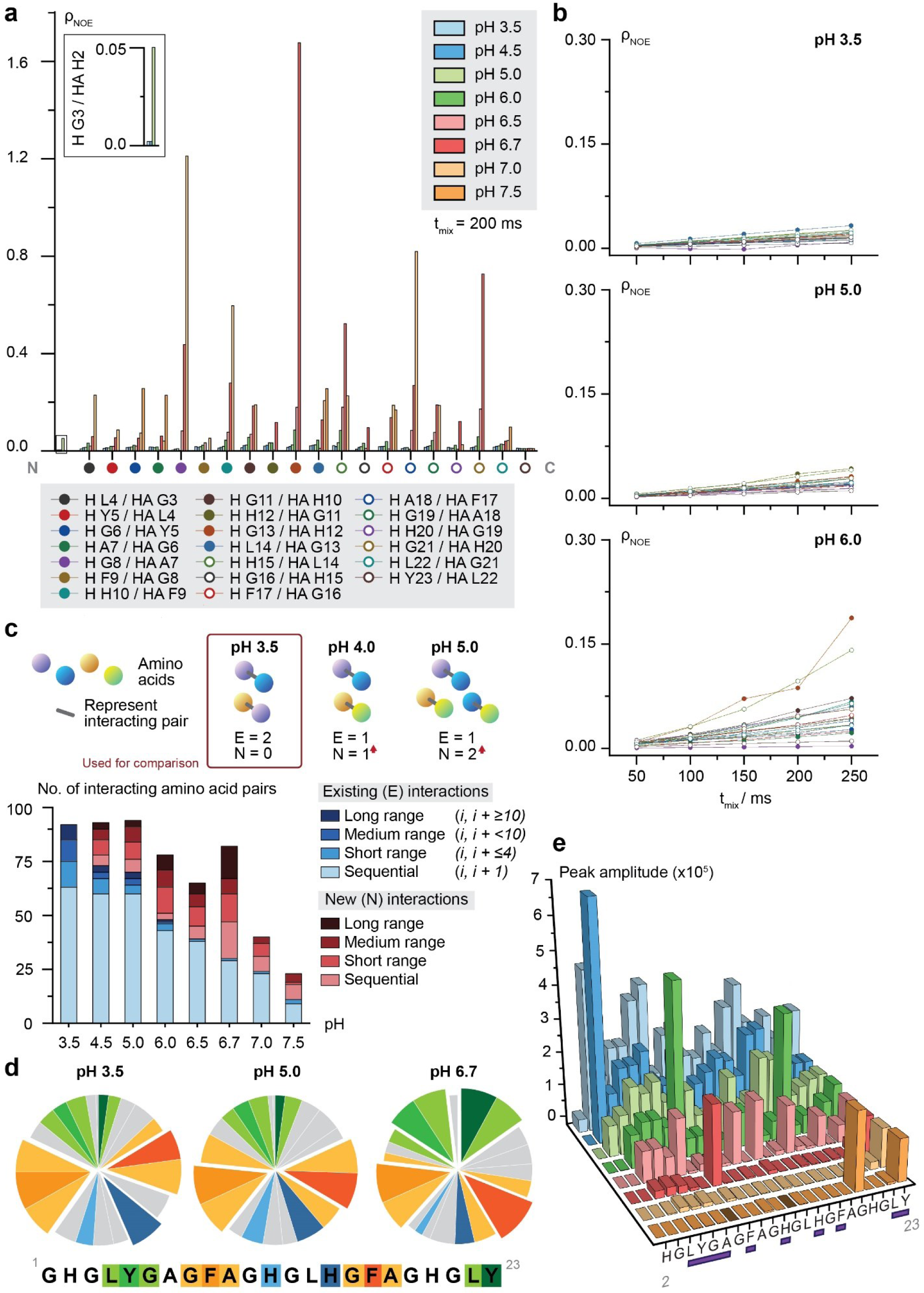
Tracked NOEs and interactions for GY23. **(a)** Bar plot of tracked NOE intensities across different pH values. Only HN/Hα (represented above as H / HA) sequential cross peaks are plotted for clarity. Missing bars indicate broadened resonances beyond detection at the particular pH(s). The inset shows a zoomed-in version of the H G3 / HA H2 bar plot on the same axes. NOE intensity, ρ, is derived from the ratio of the cross peak amplitude to the corresponding diagonal peak amplitude with values taken from CARA software^24^. **(b)** Plots of HN/Hα NOE intensities against mixing time at different pH for GY23 only. Lines are included for visual aid. **(c)** Classification of tracked inter amino acid interactions into different types at each pH. Interactions are further identified as existing or new ones based on interactions at pH 3.5. A graphical summary of the computation method is in the legend. **(d)** Classification of tracked inter amino acid interactions based on residues. Each slice represents one amino acid residue in GY23 (except for G1 and H2 – often broadened beyond detection). Slices are coloured according to in-figure legend, with the rest in grey. Pie charts at other pH conditions are in Fig. S5. Statistically significant slices are offset to highlight corresponding residues. **(e)** Plot of HN/HN diagonal peak for each amino acid of GY23, except for the first residue, glycine. Values for the peak intensities were determined using CARA after ensuring consistent spectral processing. Bars are coloured according to the different measurement conditions as indicated in the legend of panel (a). Only pH values are indicated. Buffers were prepared as detailed in the Methods section.

### Peptide-specific variations in NOE intensities

For GY23, the NOE intensities markedly increased near neutral pH (Fig. 2a), and the increase was less pronounced at the peptide termini, likely due to their conformational flexibility. For (GY23)_2_ and GW26, similar increases in NOEs were evident at higher pH. However, their more robust phase separation propensities led to challenges in assigning and tracking unambiguous NOEs due to significant peak broadening, particularly under conditions beyond pH 6.0 (Fig. S4). Notably, (GY23)_2_ exhibited greater NOE intensities compared to GY23 and GW26, possibly due to its larger MW. In contrast, despite GW26’s higher phase-separating propensity, the NOE intensities were surprisingly comparable to those of GY23 (weaker propensity) across all conditions, although the increase in NOE intensities from pH 3.5 to 6.0 was more gradual for GW26. This could be due to the different nature of these two HB*peps,* where GY23 has hydrophobic motifs favouring gel-like structures, distinct from GW26, which is comprised of charged linkers resulting in more liquid-like coacervates^23^. Hence, GW26’s greater ability to phase-separate is counteracted with a more liquid-like behaviour yielding an increase in NOE intensities for GW26 smaller than expected.

### New interactions observed across varying pH

Structural interpretation of TrNOEs required filtering off the NOEs present in the free peptides in solution. For the structural analysis, we selected TrNOEs manifesting as a 3-fold additional intensity of NOEs in acidic solution, or as new NOEs absent in the low pH spectra (Fig. S6). As the sticker and spacer residues of GY23 are well characterized^18,25^, along with a more complete dataset extending to pH 7.5, we focused on GY23 and, by relation, (GY23)_2_ for subsequent analysis. Using the collective interactions at pH 3.5 as a base, we found that the number of new interactions increased with pH until pH 7.0. Most new interactions formed at pH 6.7, featuring a significant number of long-range interactions. Initially, the key residues in GY23 involved in inter-amino acid interactions (Fig. 2d & Fig. S5) centred around H15 and the two Phe residues (F9 and F17) at low pH. However, at higher pH levels (6.5 and 6.7), these interactions predominantly involved the Tyr residues Y5 and Y23, underscoring their role in driving phase separation.

### Identification and stabilization of interaction nodes, role of Phe residues, and solvent exchange protection of amide protons

New TrNOEs at pH 6.7 were analyzed and categorized into groups of supported interactions to identify interaction nodes (Fig. 3b, dashed lines). These nodes matched previously proposed nodes based on mutagenesis studies (Fig. S7). Calculated structural conformers suggested that Tyr residues were involved in π-π stacking with each other as well as in hydrogen bonds with His residues, contributing to peptide stabilization within droplets (Fig. 3d & Fig. S8). Neighbouring residues like Leu were found to stabilize the interactions. This was further confirmed with quantum chemistry calculations (Fig. S9a, b) and experimentally since a GY23 variant lacking L4 showed impaired phase separation and formed aggregates despite the presence of key residues (Fig. S9c).

**Figure 3.**
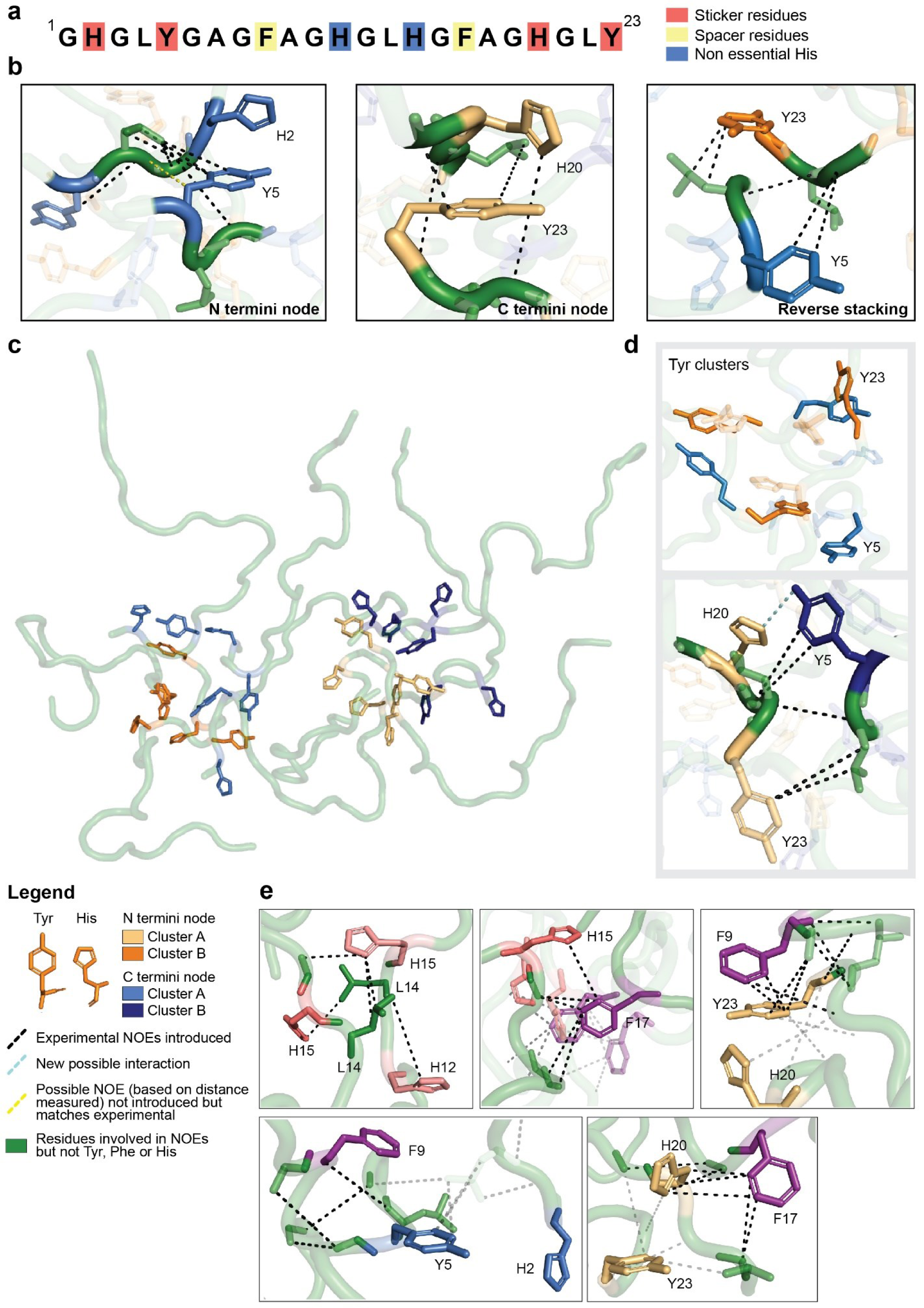
Snapshots of calculated structures of interacting peptides (GY23) from new inter amino acid TrNOEs obtained at pH 6.7. **(a)** Sticker and spacer residues of GY23 as identified in past studies, along with highlighting the two middle (in sequence) His residues with previously undefined roles. **(b)** Depiction of key residues in interaction nodes (schematic as shown in Fig S8b) with experimental NOEs shown as black dashed lines. **(c)** Arrangement of 10 peptides with experimental NOEs incorporated. The corresponding schematic can be found in Fig. S7c. Key His and Tyr residues are rendered in stick representations and coloured according to the in-figure legend. **(d)** Depiction of Tyr clusters present from the calculated structures and additional interaction between key residues, also derived from calculated structures. **(e)** Depiction of other interactions from experimental NOEs beyond those in (b). For H15 and F17, different structural conformers are shown with reduced opacity to highlight the dynamic characteristics of such interactions. Other structural conformers are not shown for other interactions for clarity purposes but are observed to be dynamic.

Other interactions along the peptide sequences, identified from TrNOEs, including the middle His and Leu with F17, likely promote stability but were noted to be dynamic across structural conformers (Fig. 3e). Isolated TrNOEs involving pairs such as H15-Y5 and F9-F17 were observed but not used for structural calculations due to insufficient additional supporting NOEs at pH 6.7.

Phe residues act as spacers in GY23, with their mutations delaying phase separation kinetics. Experimental TrNOEs revealed that Phe residues stabilized key interactions (Fig. 3e & Fig. S9b) and formed additional interaction clusters, suggesting their role in regulating oligomer formation that subsequently grows into micron-sized droplets (Fig. 2d & Fig. S5). Their interactions with Tyr, His, and Phe were noted to be (energetically) promiscuous, contributing cooperatively to structural stability (Fig. S8).

His and Tyr residues at both termini of the peptide play critical roles in phase separation (Fig. 3c). Calculated structures showed that the peptides adopted expanded conformations with both ends involved in interaction nodes, matching observations of broader amide proton (H_N_) signals with increasing pH (Fig. 2e & Fig. S10a). The extent of broadening varied, indicating different solvent exposure levels since there were different extents of hydrogen exchange. Additionally, a variable decrease in amide intensities was observed when pH increased from acidic (3.5) to neutral (Fig. 2e). This indicated varied exposure levels to bulk solvent water depending on pH. Residues with gradual decreases in amide intensities corresponded to interaction nodes that were present in dynamic clusters, the former acting as primary hubs shielding amide protons from water through the surrounding side chains (Fig. 3). This trend was observed for (GY23)_2_ and GW26, with GW26 showing more uniform broadening (Fig. S10c, d).

### Granular organization of peptide clusters revealed through Small Angle Neutron Scattering (SANS)

Full contrast (100% D_2_O) SANS data for the coacervate/dense phase at neutral pH (15 mg/mL GY23 in 100 mM phosphate buffer pH 7.4, 100% D_2_O; Fig. S11) did not give signs of any form or structure factor with correlation peaks within the *Q* range studied^26,27^ (see Section 1 of Supporting Information for experimental description). Instead, there was an upturn of scattered intensity at low *Q* following a linear behaviour on a logarithmic scale. Such behaviour is consistent with the formation of μm-large coacervate droplets (measured within a few minutes of pH adjustment). Applying a power law fit to the low *Q* region, we estimated the fractal dimension to be roughly 3 (Fig. S11, Table S1), indicating a surface fractal nature of the coacervate droplets. Thus, the solvent-coacervate interface was not sharp but rather fractal-like^28,29^.

GY23 was spiked with deuterated GY23 (dGY23) and measured under GY23 matching conditions (42% D_2_O). dGY23 was also measured under acidic, dilute conditions, keeping its concentration constant (5 mg/mL). This allowed the structural units within the coacervate to be focused on and isolated under “dilute-like” conditions (Fig. S12). Surprisingly, the SANS data (Fig. 4a) were quite similar to that measured at pH 3.5 (dilute), with a slight decrease in scattering intensity and a small upturn at low *Q*. The small upturn at low *Q* is consistent with coacervate droplet surface scattering and was visible even in the GY23 42% D_2_O data set. The slight decrease in scattering intensity could be due to minor differences in sample concentration, sedimentation, or a hidden structure factor^26,27^. The Kratky plot (Fig. 4b) displayed the typical signature of intrinsically disordered proteins, containing both a folded and a disordered region.

**Figure 4.**
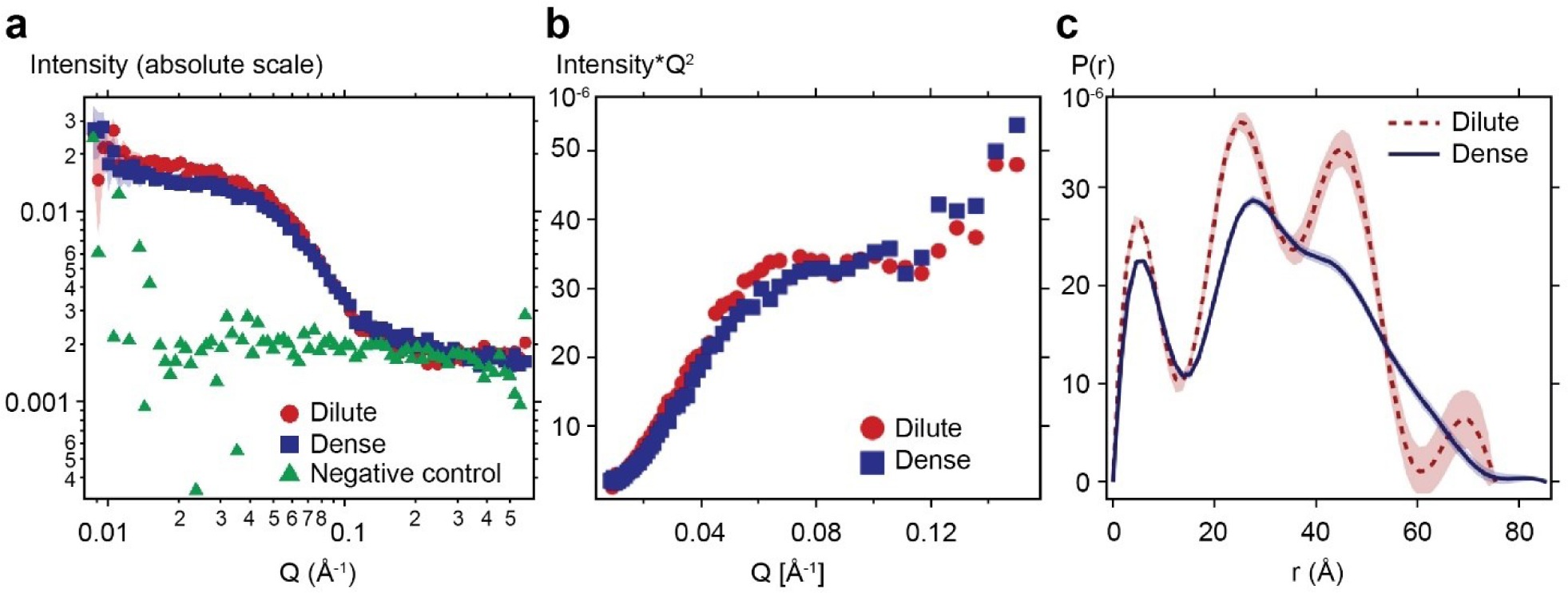
Higher order structural organization of GY23 clusters in dense phase and oligomeric association in dilute phase revealed through SANS. **(a)** SANS spectra for 5 mg/mL dGY23 in acetic acid 42% D_2_O (red circles), 15 mg/mL GY23 spiked with 5 mg/mL dGY23 in 100 mM phosphate buffer pH 7.4 42% D_2_O (blue squares), and 15 mg/mL GY23 in 100 mM phosphate buffer pH 7.4 42% D_2_O (green triangles). **(b)** Kratky plot and **(c)** pair distribution function, *p*(r), for dGY23 in dilute and dense phase.

Given the “dilute-like” conditions, a pure form factor was assumed, with *R*_g_ and *V* calculated to be 2.6 nm and 30.211 Å^3^, respectively (Table 1). The Porod volumes reported were rough estimates given the small peptide sizes studied here^30^Assuming 2723.5 Å^3^ per peptide, the structural units within the peptide cluster were calculated to consist of roughly 11 peptides each, which is close to the CYANA calculations (10 peptides in Fig. 3c).

**Table 1.** Model free SANS data analysis including radius of gyration (Rg), Porod volume and estimated number of peptides per building block.

|  | Dilute Phase | Dense Phase |
| --- | --- | --- |
| $R_g$ (guinier) ($\text{\AA}$) | $25.70 \pm 0.02$ | $26.10 \pm 0.01$ |
| Porod volume ($\text{\AA}^3$) | 12476 | 30211 |
| Number of peptides/building block | 5 | 11 |

### Small oligomers are detected at non-coacervating conditions

To probe the starting configuration of the peptides leading up to phase separation, we briefly looked at GY23 in non-coacervating conditions, *i.e*. low pH. SANS results, supported by NMR data, indicated the presence of small oligomers. The peptide ends were conformationally flexible, featuring non-canonical helical elements (mostly single 3-turns) along the peptide chain (Fig. S13b, e). Additional details can be found in Section 2 of Supporting Information.

### Estimation of the droplet surface area mediating exchange with free peptides using TrNOEs

The analytical expressions for TrNOEs in a two-spin system in the absence of ligand-protein cross relaxation^31^ were used to assess the surface area of the coacervates mediating the exchange of free and bound peptides. Calculations, performed using a reasonable set of assumptions (Section 3 of Supporting Information) demonstrated that the outer surface of dense coacervates with the radii observed via brightfield microscopy and DLS (Fig. 1b, g), in the range of 1-5 μm, was two to four orders of magnitude insufficient to produce TrNOEs exceeding a detectability threshold of 10 % of the diagonal peak intensity. Figure 5a illustrates that the TrNOE detectability limit, represented by the lower contour line, could only be achieved if coacervates radii were smaller than 0.01-0.04 µm, which was not consistent with coacervates observed by optical microscopy and DLS. This is due to the higher surface area-to-volume ratio in smaller spheres, indicating that a relatively larger proportion of the material was near the surface compared to the interior. On the other hand, coacervates with a radius of 1 µm containing numerous inner spherical cavities of 1.5 nm (Fig. 6) were calculated to provide a sufficiently extended inner interface capable of mediating the peptide exchange, resulting in TrNOEs exceeding the detectability threshold (Fig. 5b). Calculations of the detectability conditions of TrNOEs using a range of dissociation constants (*K*_D_) estimated using the phase transition diagrams in Fig. 1 and realistic diffusion-limited rate constants (*k*_D_) (Fig. S14 & S15) revealed that the particular choice of these constants within their realistic ranges could only slightly affect the detectability of TrNOEs.

**Figure 5.**
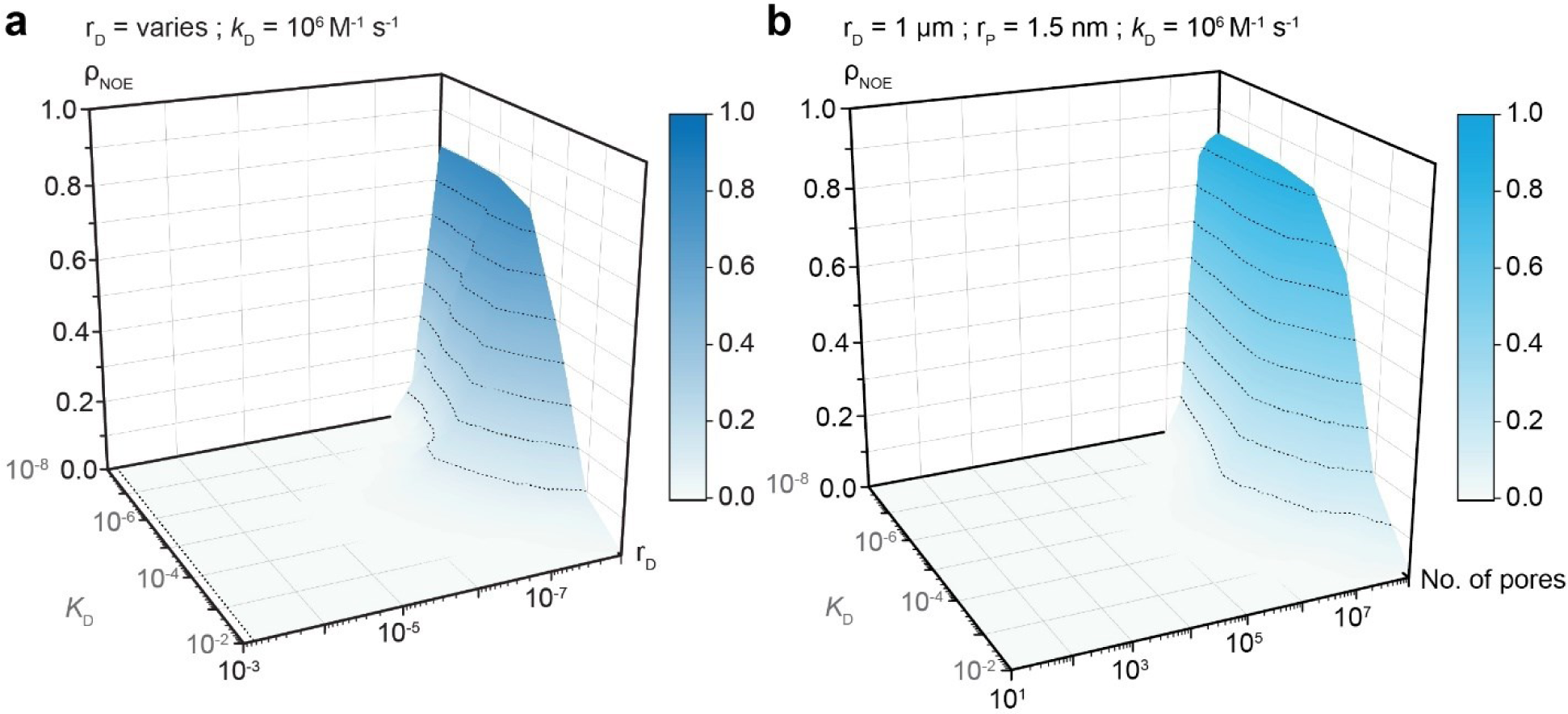
Analytical calculation of TrNOE intensities. **(a)** NOE intensities are calculated as a function of the dissociation constant, *K*_D_ and droplet radius, r_D_, with a fixed diffusion-limited rate constant at 10^6^ M^-1^ s^-1^. **(b)** NOE intensities are calculated as a function of *K*_D_ and the number of pores. The radius of the droplet, r_D_, is fixed as 1 μm, and the pore radius, r_P_, is predefined at 1.5 nm with other parameters, same as in (a). Detailed calculations and discussions of these parameters can be found in Section 3 of Supporting Information. NOE intensity, ρNOE, is defined as the ratio of the intensity of the cross peak over the intensity of the diagonal peak. Dashed lines on graph plots correspond to an interval of 0.1 on the colour scale as a guide for the eyes. Related graph plots can be found in Fig. S14 and S15.

**Figure 6.**
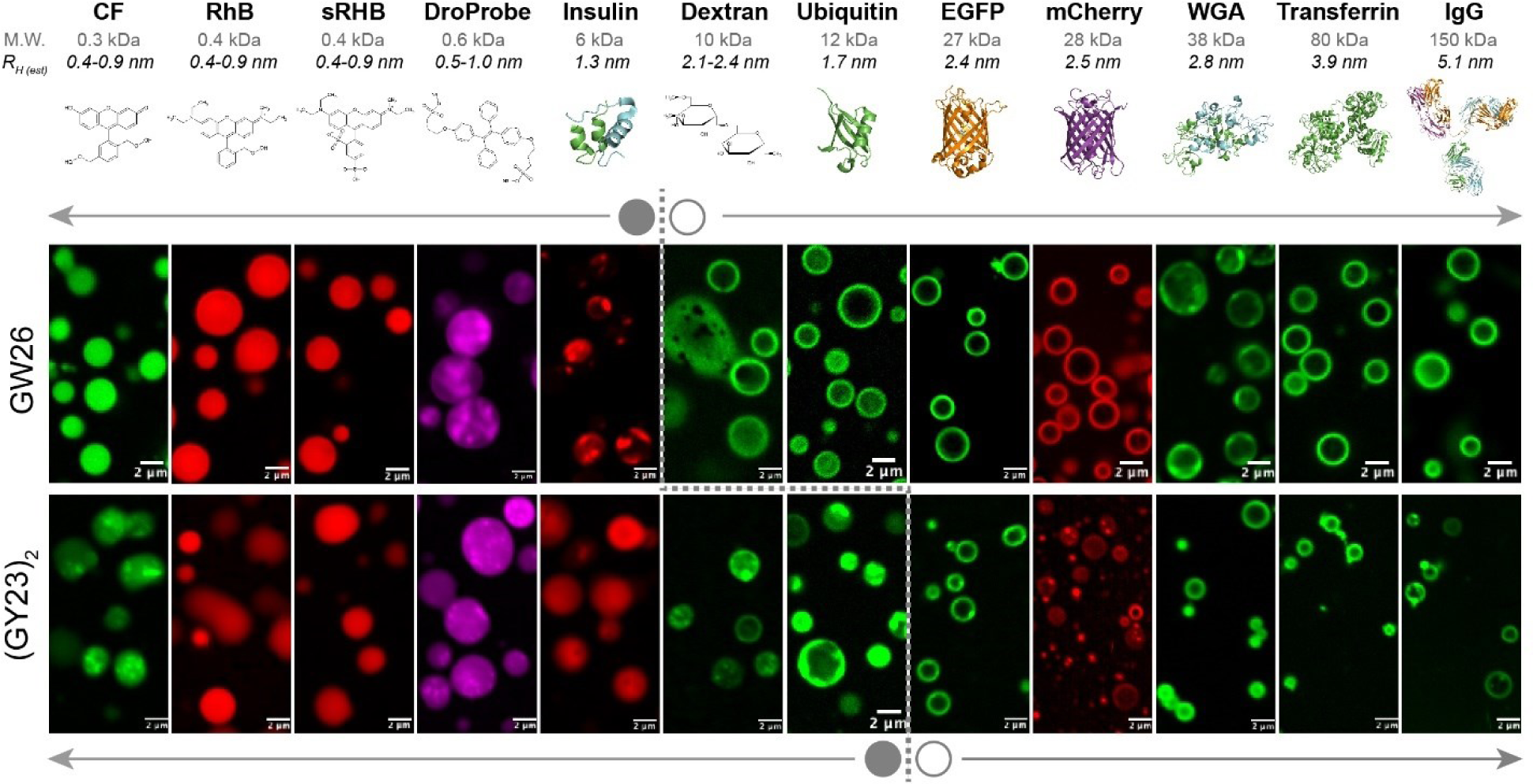
Visualization of different cargo molecules with HB*pep* variants – GW26 and (GY23)_2_. Confocal microscope images of various fluorescently labelled cargo molecules mixed with pre-formed coacervates: GW26 (top panel) and (GY23)_2_ (bottom panel). For GY23, though attempts were made, the droplets formed were too small for effective imaging of the distribution of cargo within and outside of the droplets. Scale bar indicated in the micrographs corresponds to 2 μm. Molecular weights (M.W.) of the cargo are listed, and the hydrodynamic radius (R_H_) of the molecules was estimated correspondingly from the calculator available in the Malvern Zetasizer software^32^. For small molecules, the R_H_ reported is a range estimated for linear polysaccharides, dendrimer polymers and branched polysaccharides, whereas for proteins used, the shape of the molecules is assumed to be globular. PDB structures presented are coloured according to the number of chains present.

### Estimation of cavity sizes through confocal microscopy

By incubating pre-formed coacervates with fluorescently-labelled cargo molecules ranging in MW from 0.3 kDa to 150 kDa (∼ 0.5 nm - 5 nm) and letting these cargos adsorb within the coacervates, a size cut-off was identified for them to be evenly distributed within the whole coacervate droplet: confocal images revealed a clear size-selectivity at ∼ 10 kDa. Cargo molecules above this size remained near the coacervate corona, whereas smaller cargo molecules localised within the coacervate interior. An analysis of the predicted LogP values for small molecules and surface hydrophobicity for proteins, as detailed in Table S2, suggested that the observed partitioning was mostly independent of the hydrophobicity or hydrophilicity of the cargo molecules examined. Therefore, the results presented in Fig. 6 indicated that the coacervates functioned as molecular sieves, selectively allowing cargo molecules to diffuse within the interior based on size, which is consistent with the presence of finite-sized voids that cut off larger MW cargos.

## DISCUSSION

Achieving meso-to atomic-scale resolution of structures formed within the dense phase during LLPS, typically appearing as spherical microdroplets, is challenging^33,34^. In optical microscopy, the droplet interior appears as a homogeneous phase with no distinguishable features unless illuminated by fluorogenic reporters^6,35,36^. Yet, a higher structural resolution remains elusive. Only EM-based methods have been able to identify mesoscale structures, like convoluted cavities and pores, within droplets involving relatively large proteins^1–3^. It is not yet clear whether these inner structures are unique only to the respective systems or if they are universal features appearing in a broad range of coacervates, including those formed by small peptides and other molecules.

Here, we show that a consorted application of TrNOESY, SANS and confocal microscopy to three phase-separating peptide systems reveals a hierarchical structural organization within the coacervate dense phase spawning the specifically structured clusters of the sticker and supporting residues organized into well-defined oligomers, which in turn associate with each other to form mesoscale order in droplets. The observable and notable increase in TrNOEs tracked from the solution phase (at pH 3.5) to the phase-separated conditions near neutral pH is fundamental as it highlights the dynamic but topologically persistent structures in NMR chemical shifts timescales. A conservative analysis of TrNOEs detectability in coacervates, using realistic free-to-bound peptide equilibrium and exchange reaction rates, indicates that TrNOEs are not detectable for dense and homogenous coacervates due to the insufficient amount of peptide exchangeable sites on the coacervates’ surface as depicted in Case 2 of Fig. 7. Our analysis implies that the coacervates should have significant voids organized into a porous network, which may result from the loose association of smaller peptide oligomers (Fig. 7, lower panel). TrNOEs are thus observable due to the extended available surface for exchange, as shown in Case 3 (Fig. 7). These clusters of peptide oligomers inferred by NMR are directly observed using selective deuteration and SANS. These observations are also corroborated by recent literature on spatial inhomogeneity within droplets^6,7^ and by previous studies indicating the presence of voids^1–3,37^ as well as pores in GW26 coacervates uptaken by mammalian cells^38^.

**Figure 7.**
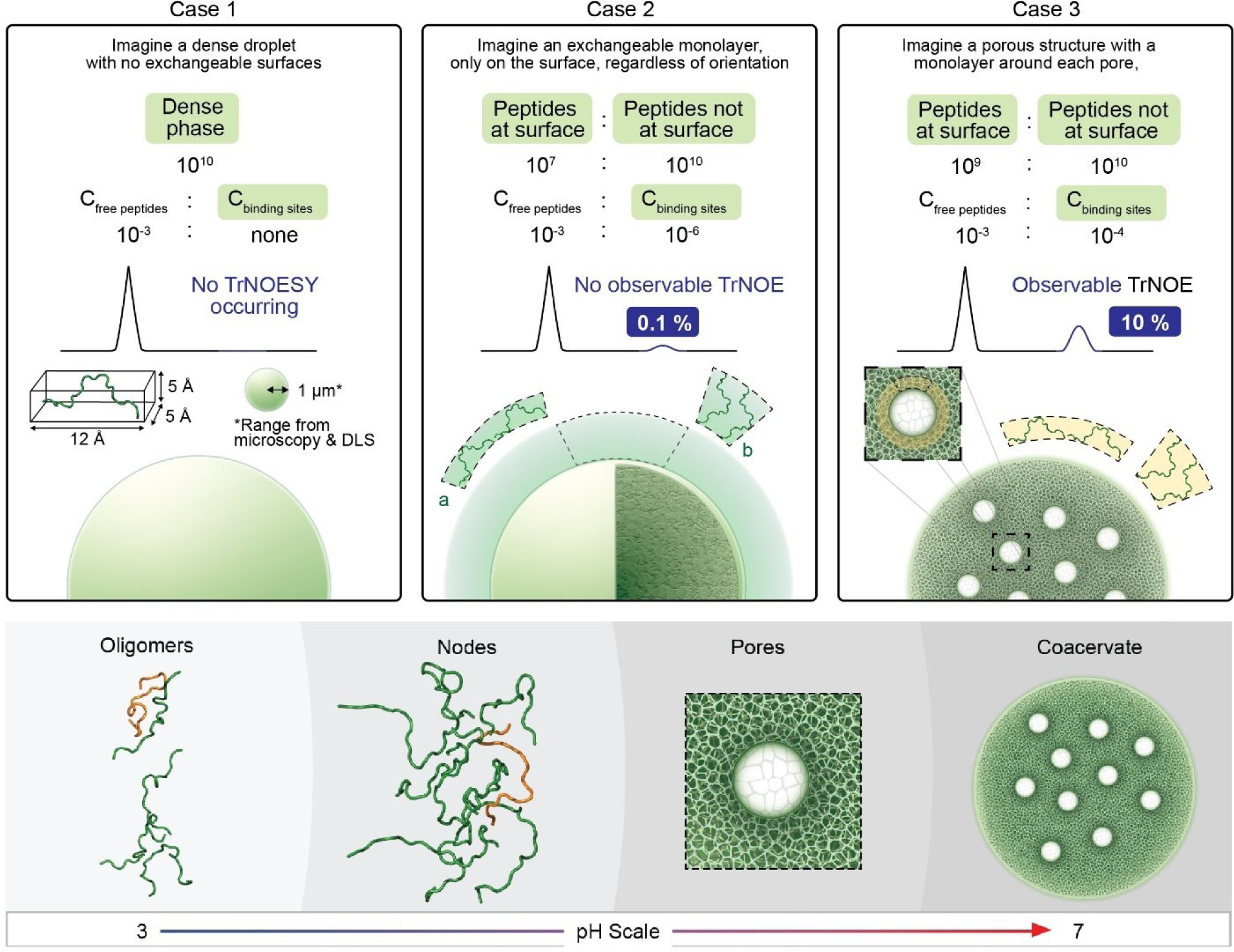
Schematic illustration of exchanging regions and transition from small oligomers to coacervates. **Upper panel:** Different cases are depicted where the exchanging regions may be to give rise to observable TrNOEs. Detailed calculations can be found in Table S3. Two possible orientations of peptides at the surface are illustrated in Case 2 (a and b). It is noted in Table S3 that the orientation did not significantly change the calculated values. The derived percentages of increase in TrNOE intensity are modulated by other system parameters, such as the dissociation constants and diffusion-limited rate constants, which in turn can influence the degree of porosity in the coacervate. However, the coacervate must still have a significant proportion of voids to give rise to observable TrNOEs (Case 3). **Lower panel:** The hierarchical structural organization of peptides within the coacervates is depicted. During phase separation induced by pH, small, initial oligomers interact through interaction nodes to form larger oligomeric clusters, which further associate dynamically into a porous mesoscale network.

At low pH, the peptides were found to form small oligomers not visible by optical microscopy. This was confirmed by both SANS, the translational diffusion coefficients derived from DOSY NMR and by NOESY. Our results indicate that these small oligomers containing 5-7 peptides may represent the initial nucleation clusters for macro phase separation. Subsequently, as pH increases to around neutral pH, these small oligomers may associate to form larger clusters, which in the dense phase contain 10-11 peptides. These clusters are driven mainly by π-π stacking between Tyr residues located at the peptide termini, which brings the sticker residues into close proximity.

Although we currently lack direct experimental data to estimate the fraction of all possible stabilizing interactions that are fully satisfied at any given timepoint, our cumulative observations strongly suggest that these interactions are essentially dynamic. They persist within the timescale of NMR chemical shifts and reform over longer timescales. This suggests that while these coacervates remain fluid, the existence of interaction nodes, underpinned by the formation and association of clusters, confers the role of “stickers” onto key residues mediating associations between clusters. Through the exchange of material (peptides) between clusters and non-persistent interaction nodes, these clusters connect in a dynamically rearranging granular network, where the spaces between them contribute to the droplet’s porosity. Remarkably, aside from pairwise interactions, the detected TrNOEs also highlight the molecular contributions of stabilizing residues. Many of the sticker interactions also involve neighbouring residues. Although such interactions may be relatively weaker than direct pair-wise interactions, stickers do not always interact in an optimized geometric configuration, making these additional interactions important collective contributors.

Strikingly, despite having almost the same amino acid composition, (GY23)_2_ is able to phase separate at a much lower peptide concentration (around 30-fold less) as compared to GY23 and has a closer phase separation behaviour to GW26. Structural topology (*i.e*. formation of the interaction nodes) is expected to be preserved at the peptide termini. However, the sticker residues located in the middle of the sequence most likely form more dynamic networks of interactions and serve as branching points for other peptides to connect and form a volume-spanning network, similar to the multiple Tyr residues found in GW26. This was tested in one (GY23)_2_ variant (data not shown) whereby the middle Tyr residues were mutated to Ala: although the required peptide concentration increased, it was still significantly lower than for GY23. As such, (GY23)_2_ can be thought of as a longer linker tethering two interaction nodes as compared to GY23, with higher valency. Effective spacing between the interaction nodes is, therefore, critical in driving phase separation by balancing the steric clashes and maintaining an optimal density of entangled chains to achieve structural integrity of the coacervates^39^. Compared to GY23, chain flexibility may also play a role for (GY23)_2_ as its length may be able to sample more binding conformations^40^.

Finally, several recent studies indicate that peptides and proteins remain conformationally flexible after sequestration into the dense phase^41–45^. Given previous data^18^, we did not expect GY23 to undergo substantial structural changes or maintain a persistent secondary structure. However, our results highlight the utility of the TrNOESY experiment for LLPS, where subtle changes in structural propensities can be detected via NMR, along with the types of interactions that enhance our understanding of phase separation at the atomic and molecular levels. The inherent dynamic exchange of material between phases, a critical characteristic of LLPS^46,47^, provides a suitable environment for the transfer of resonances. This study demonstrates that physical separation of the dense and dilute phases is not required to apply this technique, allowing the coacervates to be studied in their native state. While different mixing times need to be tested for various material exchange rates, we applied this technique directly to our biphasic sample, GY23, without additional perturbations that could introduce artefacts into the droplet structure. Therefore, applying this technique will benefit the structural exploration of other phase-separating systems, provided that the components of interest in the dilute phase are amenable to NMR. Moreover, spiking hydrogenated peptides with deuterated peptides in 42% D_2_O resulted in a strong and unique method to isolate the structure-building signature in simple coacervates. Several studies have used SANS to establish higher-order structures within phase separating systems^8,48^. Hence, such a method, typically used for protein complexes or mixtures^49^, could also help to probe smaller oligomeric scales, providing a more comprehensive view of the molecular scale when combined with other techniques such as NMR.

The uncovered hierarchical structural organization of coacervates implies that besides the molecular rules, it is essential to consider the physical interactions governing coacervate formation. What dictates the optimal mesh size and cross-linking density? Can it be altered by external factors? Undoubtedly, multivalency is crucial for enabling the plasticity of network formation to support function while maintaining the required cross-linking density for structural stability. Additionally, with a porous interior, how does this impact the material properties of the coacervate, and what defines the interface between the dilute and dense phases? Future structural investigations should include these physical considerations, particularly in understanding cargo recruitment within coacervates.

## METHODS

### Peptide synthesis and purification

All peptide variants used in this study, except for GW26, were prepared as described in our earlier study^25^. Briefly, peptides are synthesised on an automated microwave peptide synthesiser (Liberty Blue HT12^TM^). Subsequent purification through reverse phase High-Performance Liquid Chromatography (HPLC) with a Zorbax Stable Bond 300 C18 column from Agilent using a linear gradient of 10 % - 90 % solvent B over 60 min was done. Buffers used for HPLC were A: MilliQ + 0.1 % TFA and B: acetonitrile + 0.1 % TFA. Desired fractions were identified through mass spectrometry, pooled, and lyophilised to remove acetonitrile. GW26 was purchased from GL Biochem (Shanghai) Ltd., China, with > 95 % purity. No further purification was performed, and the powder was solubilised directly in 10 mM acetic acid (pH 3.5) to obtain the working stock. Sequences of the peptides can be found in Fig. 1a.

### Peptide and buffer preparation

The lyophilised powder of the peptides obtained was solubilised with 10 mM acetic acid (pH 3.5). Fresh buffers were prepared, ranging from pH 4.5 to 7.5, as below. Coacervation was induced by mixing peptide and buffer in a 1:10 ratio to achieve the desired final working concentrations. Buffers are prepared as per Table S4 with 0.1 M, 0.5 M, and 1.0 M ionic strength (adjusted with NaCl). Ionic strength is determined as the sum of the molar concentrations of the buffer salt and NaCl.

### Confocal microscopy of cargo molecules and HB*peps*

5(6)-Carboxyfluorescein, Rhodamine B, Sulforhodamine B sodium salt, (Human) Insulin-FITC labelled, Fluorescein isothiocyanate–dextran (avg. mol wt 10,000) were purchased from Sigma Aldrich (Singapore). DroProbe (Sodium 1,2-bis[4-(3-sulfonatopropoxyl)phenyl]-1,2-diphenylethene, BSPOTPE) was purchased from AIEgen Biotech Co. Ltd. (Hong Kong). Transferrin (from Human serum) and wheat germ agglutinin (WGA), both labelled with Alexa Fluor 488, were purchased from Invitrogen, ThermoFisher Scientific (Singapore). IgG from human serum (I4506) was purchased from Merck (Singapore) and was labelled with Alexa Fluor 488 NHS ester purchased from Thermo Fisher Scientific (Singapore) by following the manufacturer’s protocol. The free dye was removed by PD-10 desalting columns purchased from Cytiva. Recombinant EGFP and mCherry were expressed in *E. coli* and purified in-house using standard protocols. Recombinant ubiquitin was obtained as a gift from another lab group and labelled with FITC following the manufacturer’s protocol. The free dye was similarly removed by the PD-10 desalting column. The molecular weight of ubiquitin, as reported in Fig. 6, is inclusive of its affinity tag (His tag and thrombin cleavage site) and calculated linked FITC molecules, though these additions were found not to alter the partitioning of ubiquitin into the peptide systems (data not shown).

Peptide samples for imaging were prepared by first diluting 5 µL of a 10x PBS stock solution with 40 µL Milli-Q water followed by the addition of 5 µL of the peptide stock (prepared in 10 mM acetic acid). Immediately upon peptide addition in the previous step, the samples were gently pipetted up and down at least ten times to achieve uniform turbidity. Two minutes after peptide addition, 1 µL of the cargo (small MW fluorophores, labelled proteins or labelled dextran at ∼ 1mg/mL stock concentration in PBS) was added to the sample and gently pipetted a few times. The final volume of the above sample was 50 µL with a final peptide concentration of ∼ 200 µM. Fifteen microliters of this peptide sample was pipetted as a droplet on to the cover glass of a MatTek (Ashland, MA, USA) glass bottom dish (cover glass No. 1.5). The liquid droplet was imaged using a Carl Zeiss LSM 780 laser scanning confocal microscope (inverted) with fast spectral detection (32-GaAsP array; plan-apochromat 63x/1.4 oil, FWD 0.19 mm objective; motorized stage). All images were acquired using Zen Lite software and processed using ImageJ2 (version 2.3.0/1.53f).

### Brightfield optical microscopy

Phase separation behaviour of the variants was observed using the Zeiss Live Cell Observer system with an Axio Observer.Z1 microscope body, fitted with the following objectives - Plan-Apochromat 63x/1.40 Oil and Plan-Apochromat 100x/1.4 Oil. Images were taken with a Zeiss AxioCam ICc 1 under the control of microscope software ZEN lite. All samples were prepared fresh just before examination under the microscope. For GW26, an incubation period of 30 minutes was observed after mixing before examination to allow the equilibration of peptide and buffers.

### Solution-state NMR Spectroscopy

All peptides and buffers used for measurement were prepared accordingly, as above. For pH 4.5 and above, only buffers with 0.1 M ionic strength were used. 10 % D_2_O was added to the mixture before the NMR experiments. An external reference of Sodium trimethylsilylpropanesulfonate (DSS) was used to calibrate the data obtained. The following experiments were done for this paper – natural abundance ^1^H-^1^H 2D Total Correlation Spectroscopy (TOCSY) for resonance assignment of individual spin systems and ^1^H-^1^H 2D Nuclear Overhauser Effect Spectroscopy (NOESY) for resonance assignment of spins close in space. Different mixing times (10 ms, 50 ms, 100 ms, 150 ms, and 200 ms) were used in NOESY experiments for all peptides at pH 3.5, pH 5.0, and pH 6.0. The peptides’ concentrations are 200 μM for GW26, 250 μM for (GY23)_2_, and 1 mM for GY23. To investigate the full pH range for GY23, 200 ms was determined to be a suitable mixing time and used for subsequent experiments at other pH values. Due to the poor spectral quality obtained at 10 ms mixing, the corresponding data for all peptides is not shown in this paper. All data was acquired using standard pulse programs from the TopSpin 3.5 repository on a 700 MHz Bruker Advance III NMR spectrometer at 298K. Spectra were processed using Topspin 3.5 and analysed using CARA software (http://cara.nmr.ch/)^24^. Subsequent downstream data processing was performed with self-written scripts in MATLAB (MATLAB and Statistics Toolbox Release 2012b, The MathWorks, Inc., Natick, Massachusetts, United States). Peptide concentrations used were the same for Diffusion Ordered Spectroscopy (DOSY) experiments. To minimize calculation artefacts, the buffer used for the reported data in Table S5 for pH 5.8 and above was 50 mM phosphate buffer at pH 5.8 titrated with 0.5 M NaOH. Before each measurement, an equilibration time of 15 mins for GY23, (GY23)_2_ and 30 mins for GW26 was observed. As for the measurement of ubiquitin (Table S6), 50 mM phosphate buffer at pH 7.4 was used with a final ubiquitin concentration of 100 μM. The ubiquitin used is 6.8 kDa with no fluorolabelling.

### Dynamic light scattering (DLS)

Measurements were performed with the peptides using a Malvern Zetasizer DLS (Malvern Instruments, Malvern, UK) at 298 K. An initial equilibration time of 30 s after insertion of the sample into the machine was observed. A disposable cuvette was used, and no mixing was done during the intervals. All samples were prepared fresh before measurement, with 30 min incubation observed for GW26, as per what was done for microscopy. Ten measurements separated by 60 s were taken, with 3 runs of 10 s each at every measurement point. Measurements were taken at a backscatter angle of 173°. Buffers and peptides used were prepared accordingly above. The peptide concentrations used were the same as those for NMR and brightfield microscopy – for GY23, 1 mM; for GW26, 200 μM; and for (GY23)_2_, 250 μM. Raw data was extracted from the accompanying software and replotted in OriginPro (Version 2020b, OriginLab Corporation, Northampton, MA, USA) to obtain the mean and standard deviation values. For all DLS plots in this paper, bar plots correspond to the mean values obtained with the standard deviation represented with error bars.

### Quantum chemistry calculations

Structures of interacting residue pairs were prepared using Avogadro^50^ and geometric optimization was subsequently performed with Gaussian09^51^ using the method and basis set, wB97XD/cc-pvdz, chosen for its reasonable accuracy and computational costs. Energy values (SCF) in Hartrees were computed and converted to kJ/mol. As for interacting configurations derived from structural calculations, geometric optimization was performed in a two-step process whereby certain atoms were frozen to preserve the integrity of the interacting configurations for the first round of optimization. These atoms were then unfrozen (while the rest of the optimized atoms were) during the second round of optimization. Single point energies were then calculated based on the optimized structure.

### Combined assignment and dynamics algorithm for NMR applications (CYANA)

Peptides with no restraints were first built using the standard CYANA nomenclature. An upper distance limit of 5 Å was introduced for all experimental NOEs regardless of peak intensity. For each iteration, 100 conformers were calculated, and the top 20 conformers were further evaluated primarily based on violation of the distance restraints. All guiding distance restraints, as explored for initial model construction^25^ in previous work, were released. Hence, all structures from CYANA presented here are purely based on experimental NOEs obtained. Only new inter amino acid interactions at pH 6.7 were considered for distance restraints with multiple CYANA iterations before assignment as intra- or inter-peptide NOEs. The software used was CYANA 3.98.5^52,53^. All models are examined and rendered using The PyMOL Molecular Graphics System, Version 2.4. 1 Schrödinger, LLC^54^.

### Small Angle Neutron Scattering (SANS)

SANS experiments in 42% D_2_O were conducted using the small and wide-angle neutron scattering instrument (TAIKAN) at the Materials and Life Science Experimental Facility (MLF) in Japan Proton Accelerator Research Complex (J-PARC)^55^. The neutron wavelength varied from 1.0 to 7.8 Å. A small-angle detector bank with a sample-to-detector distance of 5.65 m was used. The covered *Q*-range was *Q* ∼ 0.005–1.5 Å^−1^ in a single measurement. A banjo-type quartz cell with a neutron path length of 1.0 mm was utilized as an exposure sample cell. SANS measurements were performed at room temperature (approximately 25 °C.). After correcting for factors such as incident neutron distribution, detector efficiency, transmission, and background scattering, the SANS intensity was converted to absolute intensity per sample volume using a secondary glassy carbon standard^56^.

SANS experiments in 100% D_2_O were performed on D22 at the Institut Laue Langevin (France). Using a neutron wavelength of 6 Å and detector distances of 17.6 and 1.4 m, results in a Q range of 0.003 – 0.6 Å^−1^. D22 data was reduced using Grasp^57^. More details on sample preparation and data processing can be found in Section S1 of Supporting Information.

## Author Contributions

J.L., S.G. and C.B. designed and performed the experiments, respectively, for NMR, confocal imaging and SANS. J.L., S.G. and C.B. collected the data, analysed and interpreted the results. J.L. and S.G. wrote the original draft of the manuscript. Q.M.P. assisted with confocal experiments. H.B. assisted in the experimentation and data analysis related to SANS. H.I. assisted in the experimentation related to SANS. M.M. optimized the recombinant expression of HBP-GY23 for deuteration and provided the deuterated cell pellet for purification. L.P. assisted in the production of the deuterated cell pellet. M.C. conceptualized the experiments, coordinated the resources, provided supervision and made the acquisition of financial support for all SANS-related work. A.M. conceptualized the overall project, selected the systems for study, handled the project administration and made the acquisition of financial support for the overall collaborative project. K.P. coordinated the overall project, provided supervision, conceptualized the experiments, analysed the data and made the acquisition of financial support for all NMR-related work. A.M. and K.P. reviewed and edited the original draft.

## Acknowledgements

This research was funded by the Singapore Ministry of Education (MOE) through an Academic Research Fund (AcRF) Tier 3 grant (grant # MOE 2019-T3-1-012). We thank the NTU Optical Bio-Imaging Centre (NOBIC) at the Singapore Centre for Environmental Life Sciences Engineering (SCELSE), NTU for the use of the confocal microscope and Dr. Yong Hwee Foo for his help and suggestions with imaging. We thank Dr. Sun Yue and Poh Cheng Wei respectively for providing the labelled IgG and recombinant ubiquitin (for fluorolabelling) used in this study. We also thank Dr. Mu Yuguang from the School of Biological Sciences, NTU for his guidance and computational resources for the quantum chemistry calculations in this work. M.C. acknowledges financial support from the Swedish Research Council (2018-04833 and 2018-03990), Biofilm Research Center for Biointerfaces (Malmö University), Wennergren Foundation and IKUR Strategy under the collaboration agreement between Ikerbasque Foundation and Fundación Biofísica Bizkaia on behalf of the Department of Education of the Basque Government. The SANS experiment at the Materials and Life Science Experimental Facility of the J-PARC was performed under a user program (Proposal No. 2022A0301). We thank Batika Saxena for help with TOC.

