## Supplementary Information for "Hierarchical structural organization in bioinspired peptide coacervate microdroplets"

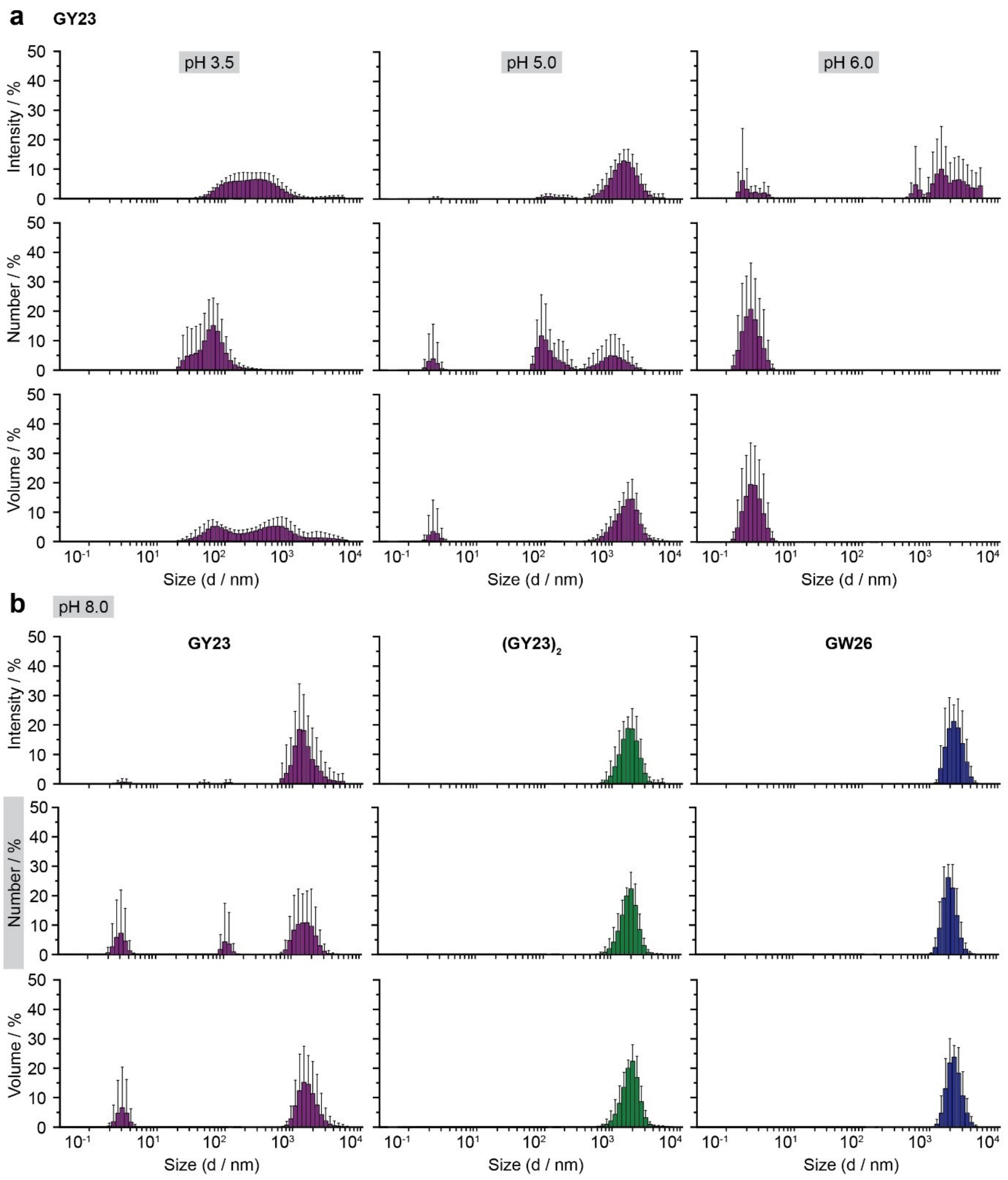


**Supplementary Figure 1. Size distribution statistical plots from DLS for HB*peps* at different pH.** Size distribution statistical plots (intensity-, number- and volume-based) obtained from DLS for **(a)** GY23 at pH 3.5, 5.0, 6.0 and **(b)** GY23, $(GY23)_2$ and GW26 at pH 8.0. The concentrations used correspond to those for NMR measurements and micrographs shown in Fig. 1(c) (main text). Details of the measurements and subsequent data analysis and plotting can be found in Methods. The number-based size distribution statistical plots for pH 8.0 have been shown in Fig. 1(g) (main text) and are included here solely for completeness. Measurements were attempted at pH 3.5, 5.0 and 6.0 for GW26 and $(GY23)_2$, but the resultant data was not acceptable and, thus, not shown.

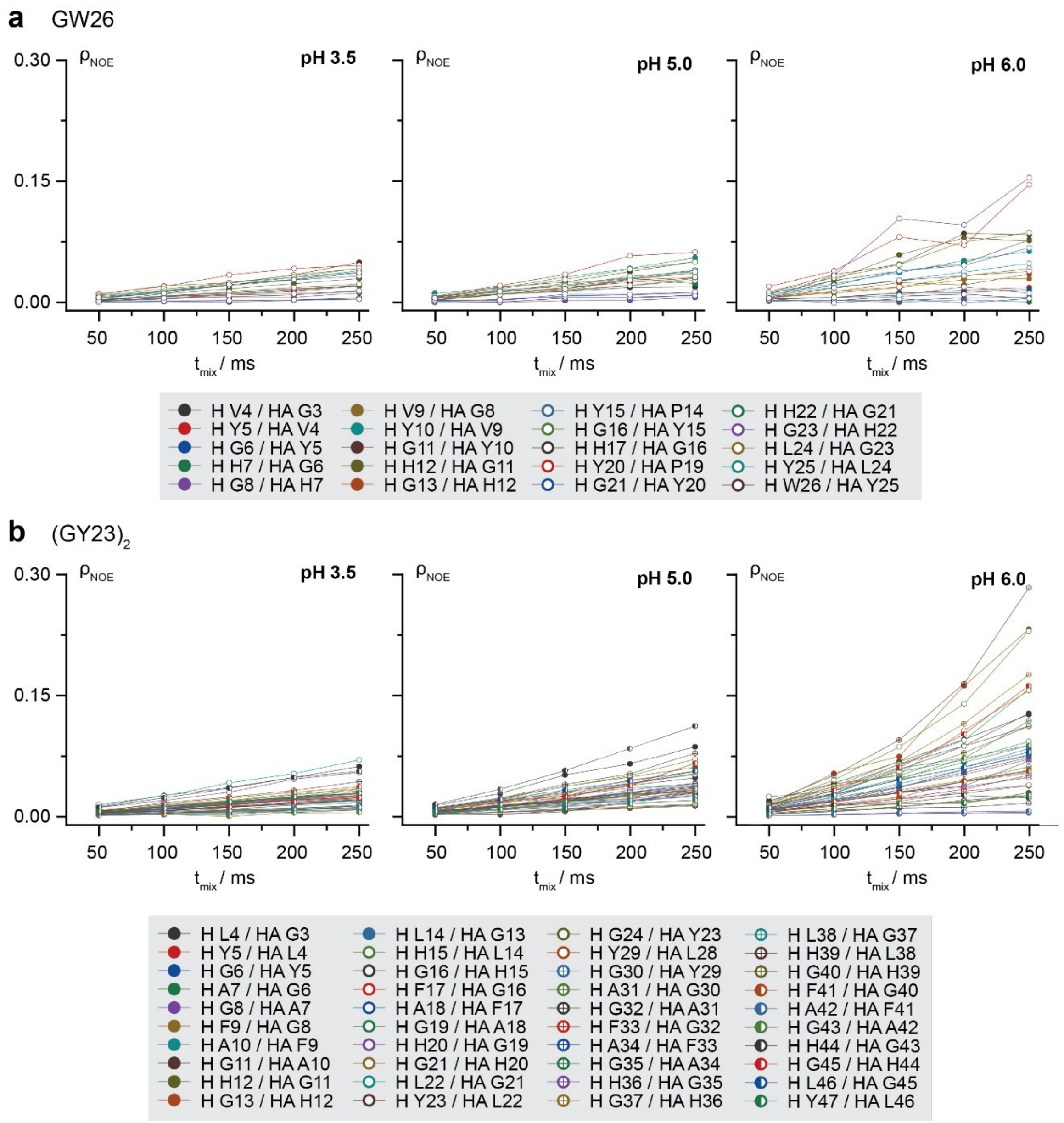


**Supplementary Figure 2. Tracked NOE intensities for HB*peps* – GW26 and $(GY23)_2$ at selected pH conditions**. Plots of NOE intensities against mixing time at different pH. Lines are drawn in to guide the eye. NOE intensity, ρ, is derived from the division of the amplitude of the cross peak by the amplitude of the corresponding diagonal peak. Peak amplitude values are taken from CARA. pH 6.0 is selected as the closest common pH across all 3 HB*peps* where NMR spectra with reasonable quality can still be obtained.

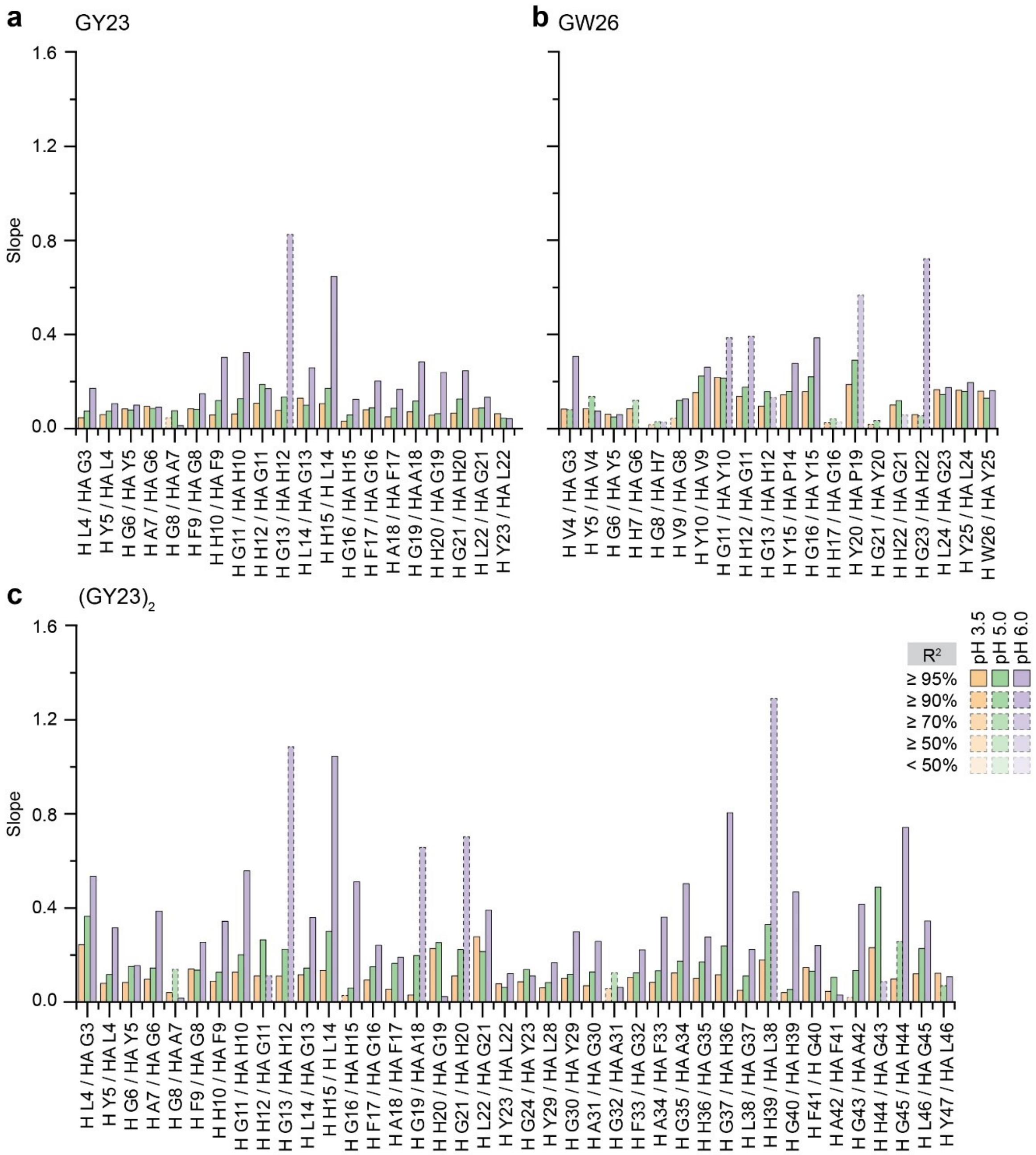


**Supplementary Figure 3. Linear regression values derived from NOE intensities for HB*peps* at selected pH conditions.** Bar plots of slope values of NOEs across different pH values for **(a)** GY23, **(b)** GW26, and **(c)** (GY23)$_2$. Only $H_N$ / $H_\alpha$ (represented above as H / HA) sequential NOEs are plotted for clarity. Missing bars indicate broadened resonances. Intensities of NOEs across different mixing times at each pH (data as shown in Figure S5 and Figure 3b) were fitted to a linear regression plot, and the derived slope values were plotted. $R^2$ values derived from the fitting are represented with different transparency levels, as indicated in the legend. Bars are also coloured according to the corresponding pH values.

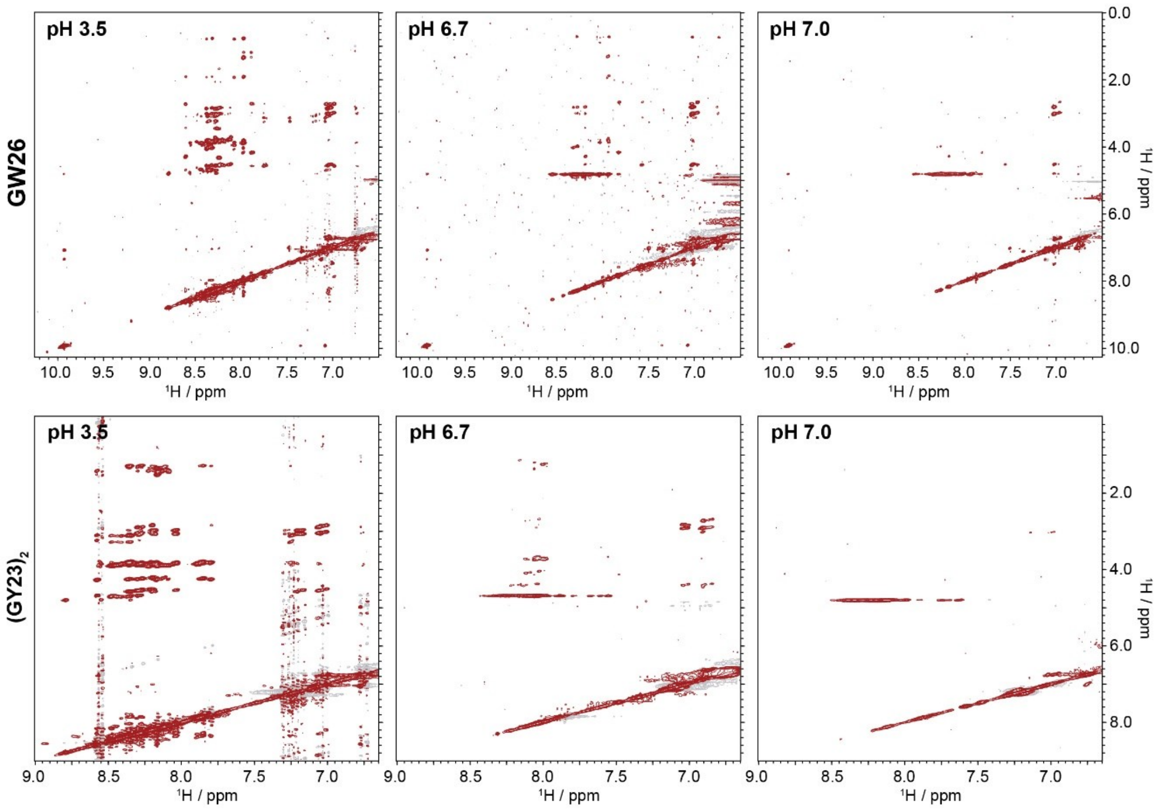


**Supplementary Figure 4. Significant broadening of peaks for GW26 and (GY23)$_2$ beyond pH 6.** 2D NMR $^1$H / $^1$H NOE plots for GW26 and (GY23)$_2$ at different pH, as indicated in the figure. Only the region corresponding to amide and aromatic protons is shown. All spectra above are processed with the same values set for threshold and contour level.

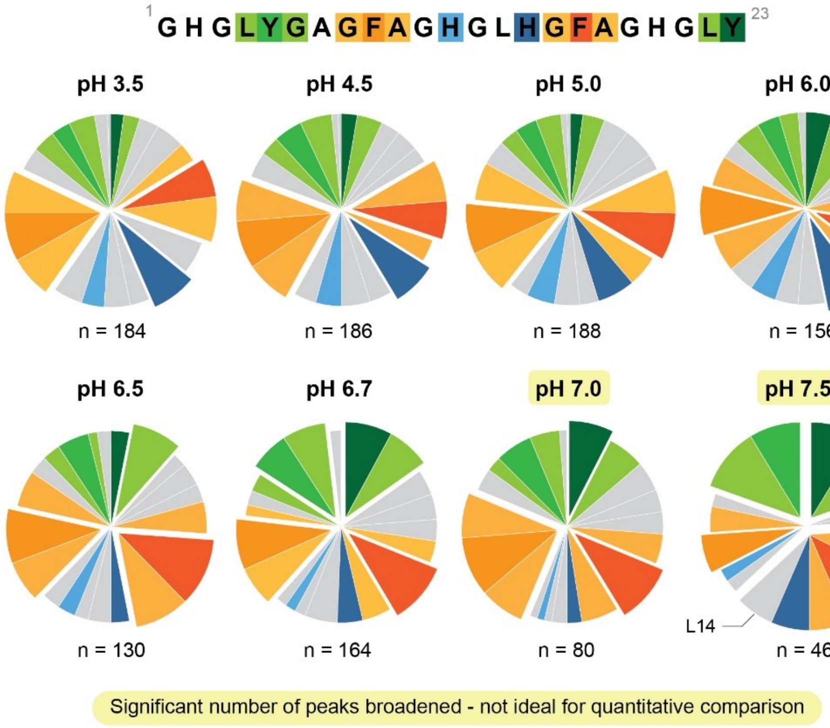


**Supplementary Figure 5. Classification of tracked inter amino acid interactions based on residues.** Each slice represents one amino acid residue in GY23 (except for G1 and H2 – often broadened beyond detection). Slices are coloured according to in-figure legend whereby rest of the amino acids are in grey. Pie charts at pH 3.5, 5.0 and 6.7 are found in Figure 2 and just presented here for completeness. Statistically significant slices are offset to highlight corresponding residues. The total number (n) of inter amino acid interactions at each condition is as shown.

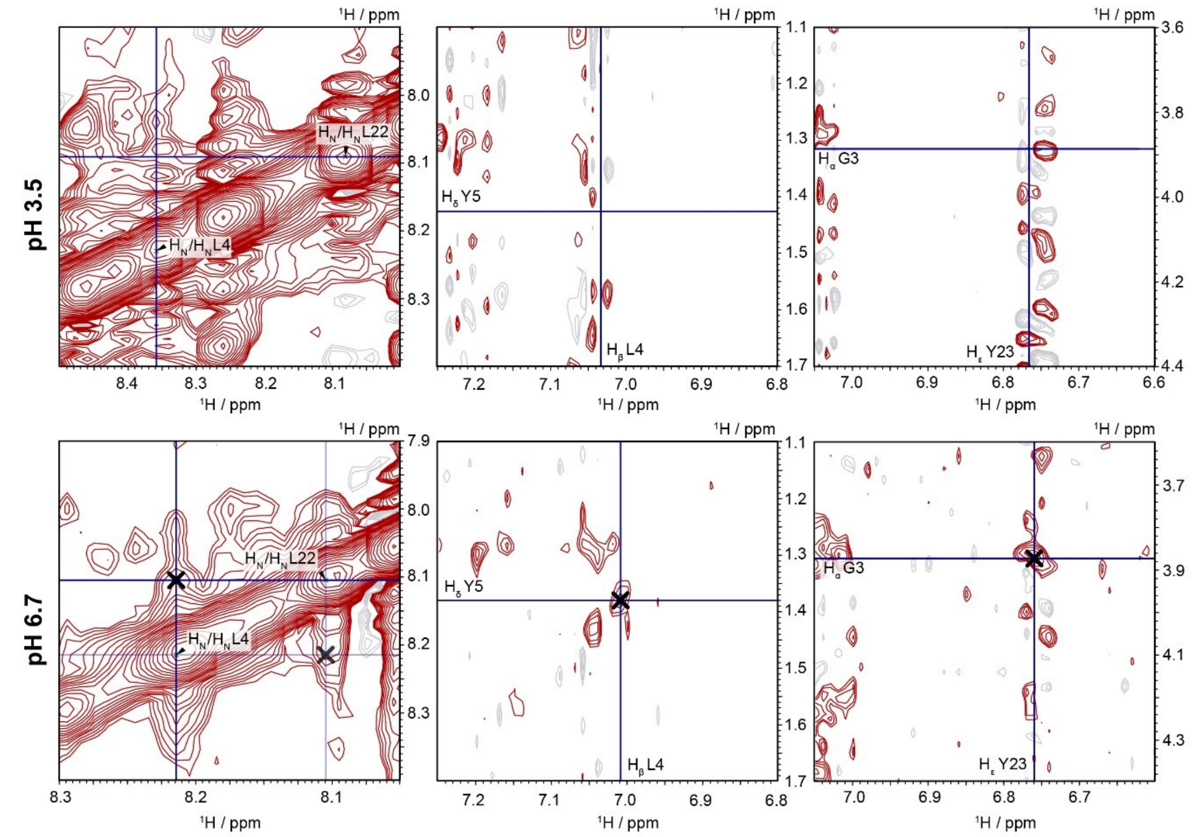


**Supplementary Figure 6. Spectra images of TrNOEs obtained at pH 6.7.** Several examples of TrNOEs obtained at pH 6.7 are depicted above with corresponding regions from pH 3.5 included for comparison. Navy lines serve as a guide for the eyes, corresponding to the vertical and horizontal ppm values of the stated protons. New peaks, *i.e.* TrNOEs, are marked with a thick black cross. All other assigned peaks have been removed for clarity.

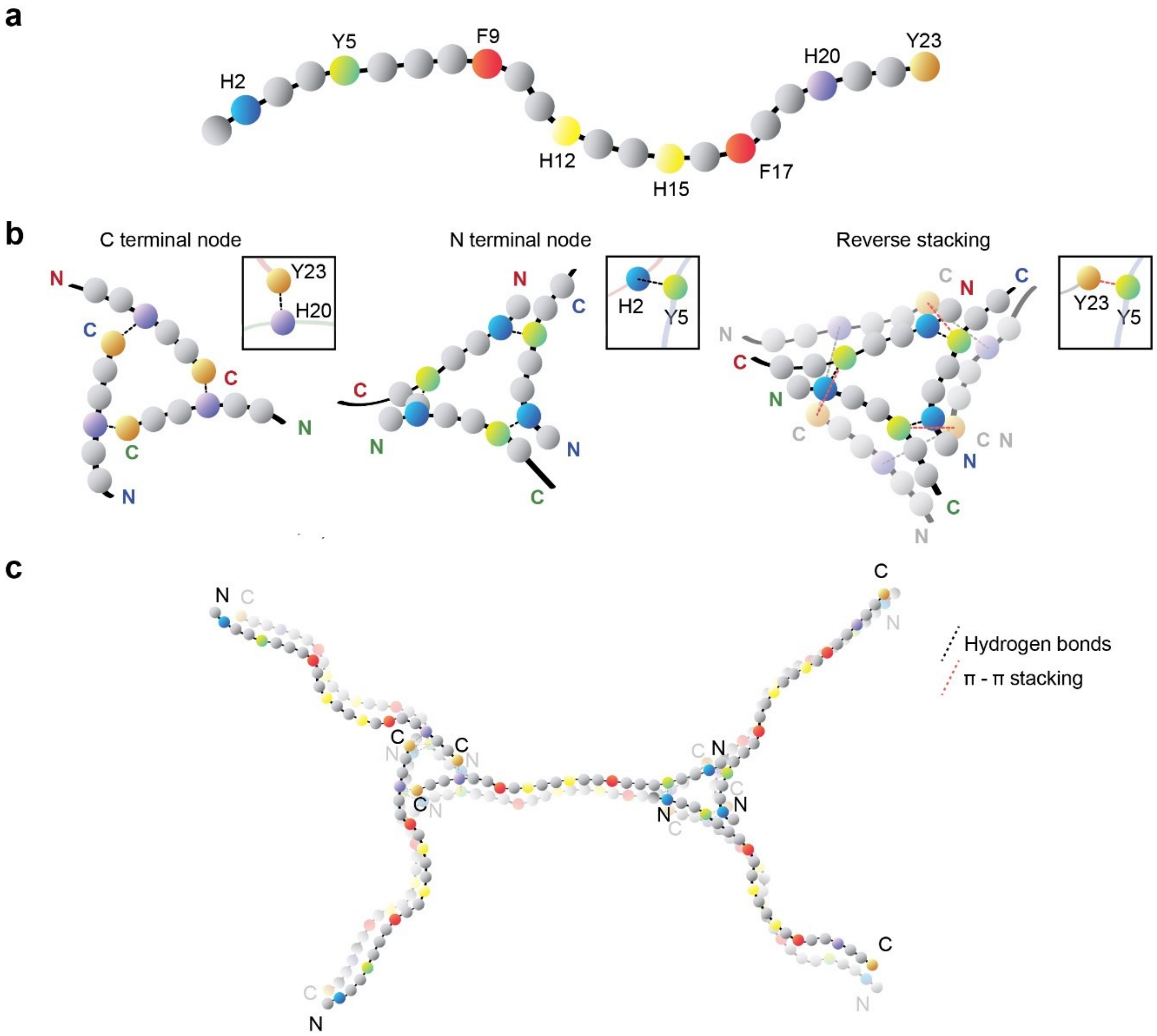


**Supplementary Figure 7. Schematic illustration of model construction for CYANA calculations. (a)** Representation of GY23 as beads on a string with important residues coloured accordingly. **(b)** Depiction of the interaction nodes at the respective terminals and stacking of the nodes with the critical residues highlighted. **(c)** Overview of the 10 peptides arranged with the interaction nodes present and terminals marked. The bottom layer of peptides has been rendered with less opacity for clarity in both (b) and (c). Adapted with permission from "Liquid–Liquid Phase Separation of Short Histidine- and Tyrosine-Rich Peptides: Sequence Specificity and Molecular Topology", by Lim, J. et al., *J. Phys. Chem. B* **125**, 6776-6790 (2021). Copyright 2021 American Chemical Society.

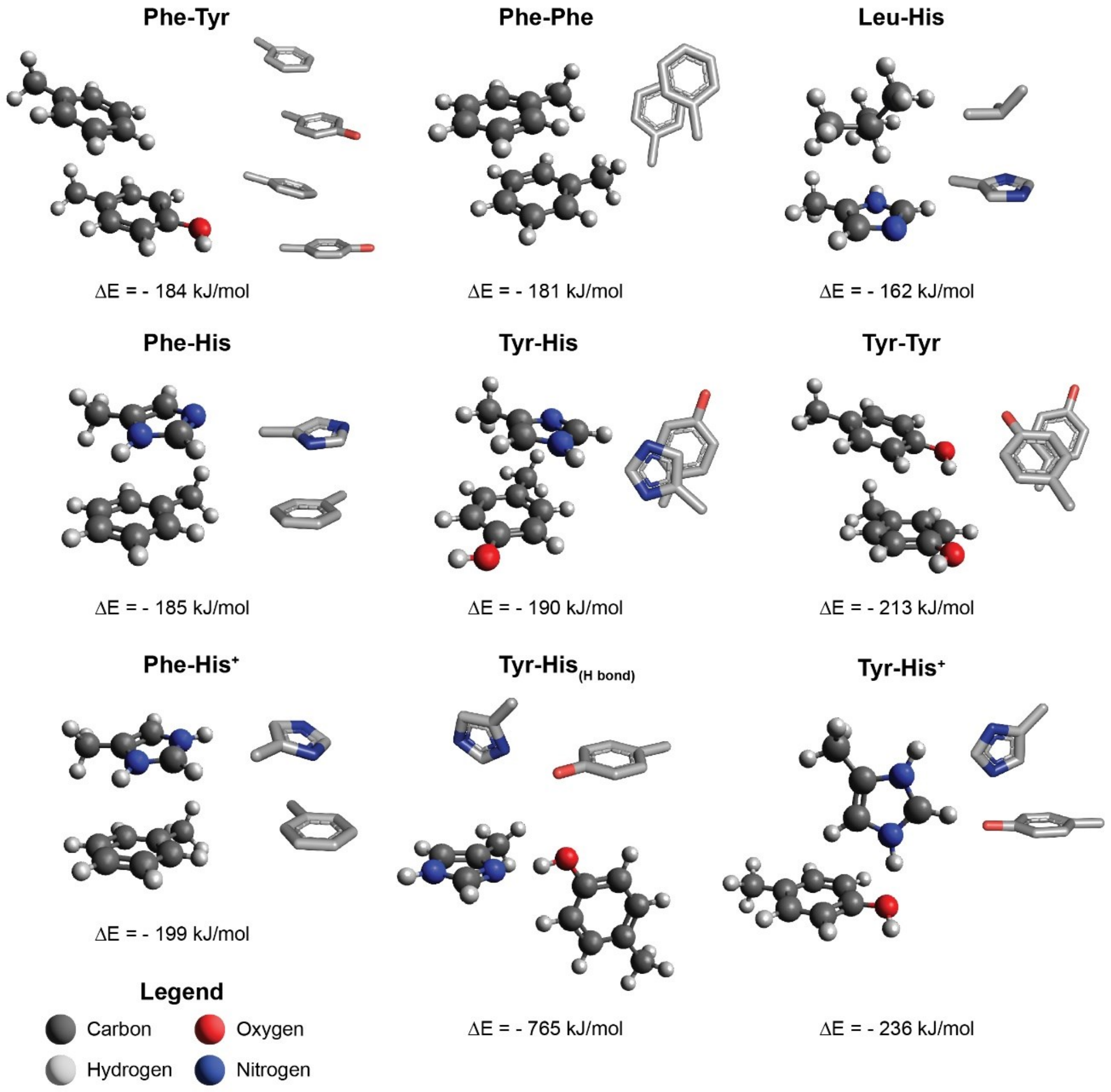


**Supplementary Figure 8. Comparison of energy values of interacting residue pairs.** Stick representations are rendered in PyMOL and coloured accordingly with different perspectives of the interacting residues. Energy values are computed, using quantum chemistry, as the difference between the single point energies (SCF) calculated when interacting residues are close together (as depicted) and the sum of individual sidechains. The above values are and should only be used for the comparison of different configurations.

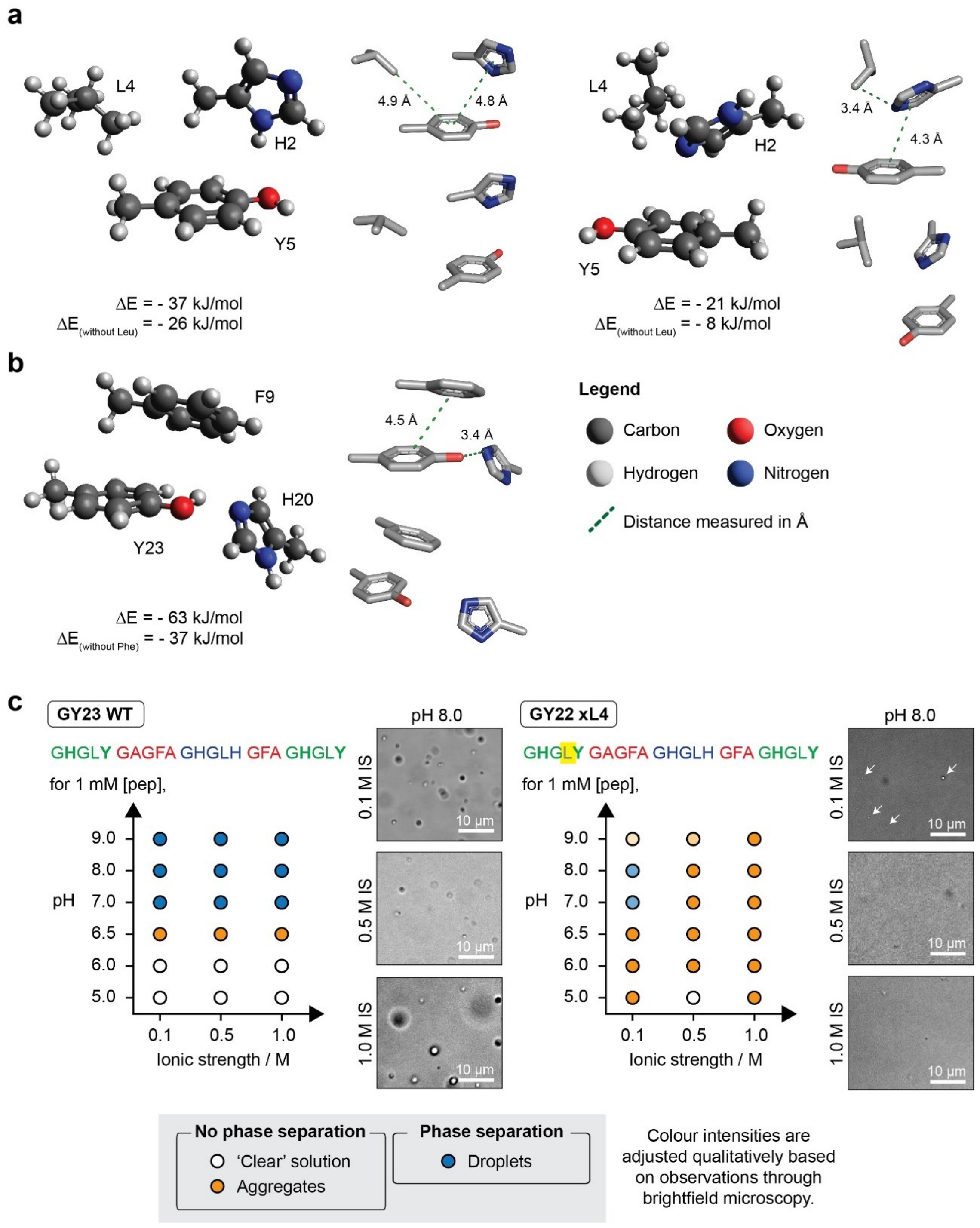


**Supplementary Figure 9. Characterization of stabilizing residues found from CYANA calculations.** **(a,b)** Structural configurations of additional residues **(a)** Leu and **(b)** Phe found with sticker residues, derived from CYANA simulations with a two-step geometric optimization (see Methods). Stick representations are rendered in PyMOL and coloured accordingly with different perspectives of the interacting residues. Distances indicated are measured using PyMOL's in-built function. Energy values are computed, using quantum chemistry, as the difference between the single point energies (SCF) calculated when interacting residues are close together (as depicted) and the sum of individual sidechains. **(c)** Plot of pH *vs* buffer ionic strength (IS) for peptide variants, GY23 and GY22 (missing L4), indicating conditions where phase separation is observed at peptide concentration = 1 mM. Corresponding micrographs taken at pH 8.0 at different IS are shown for comparison.

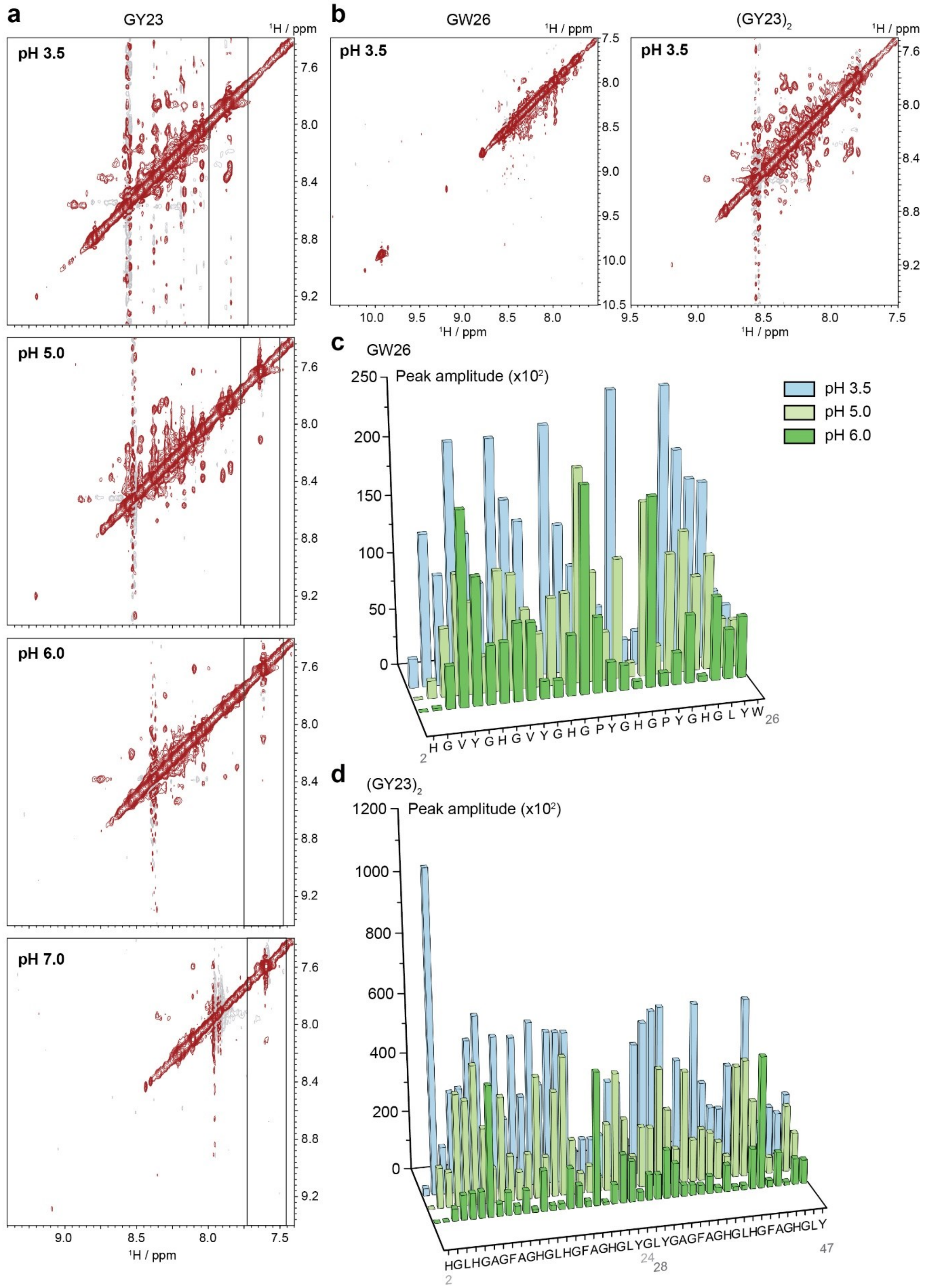


**Supplementary Figure 10. Tracking of amide proton intensities across different conditions for HB*peps*. (a)** 2D NMR $^1$H / $^1$H NOE plots for GY23 at different pH, as indicated in the figure. Only the region corresponding to amide protons is shown. For easy reference and comparison, the rectangle corresponds to the same diagonal peak across all the plots. **(b)** 2D NMR $^1$H / $^1$H NOE plots for GW26 and $(GY23)_2$ (as indicated) at pH 3.5 10 mM acetic acid.

Only the region corresponding to amide protons is shown. **(c)** Plot of $H_N$ / $H_N$ diagonal peak for each amino acid of GW26, except for the first residue, glycine. Values for the peak intensities were determined using the CARA software[1] after ensuring each spectra collected was processed similarly. The absolute values retrieved were then divided by a factor of $10^2$ for easier plotting. Bars are coloured according to the different measurement conditions as indicated in the legend. Only pH values are indicated. Buffers are prepared as detailed in the Methods section. **(d)** Plot of $H_N$ / $H_N$ cross peak for each amino acid of $(GY23)_2$, except for the first residue, glycine and Gly 25 to Gly 27 which cannot be unambiguously assigned.

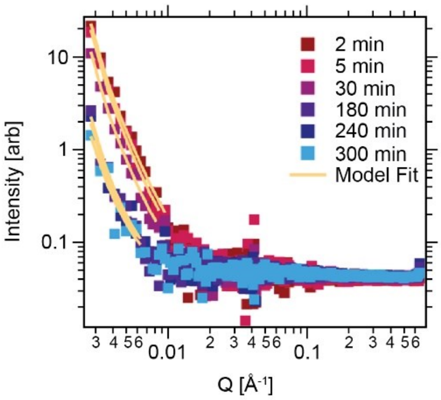


**Supplementary Figure 11.** SANS data for 15 mg/mL GY23 in 100 mM phosphate buffer pH 7.4 100% $D_2O$ over time after adjusting pH from acidic to neutral values. The overall scattering intensity increases over time during the first 30 min of measurements likely due to increasing droplet formation and droplet merging, while it decreases due to droplet sedimentation at some point between 30 min and 3 hours. Values derived from fitting the power law in the low Q region are in Table S1.

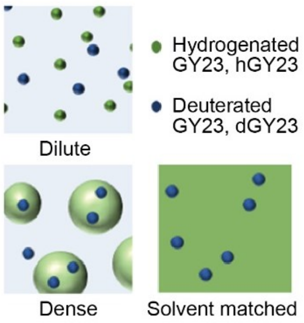


**Supplementary Figure 12.** Schematic illustrating the concept of solvent matching utilised in isolating building units of GY23 within the coacervates for SANS. The light blue background colour represents 100% $D_2O$ while the green background colour represents 42% $D_2O$.

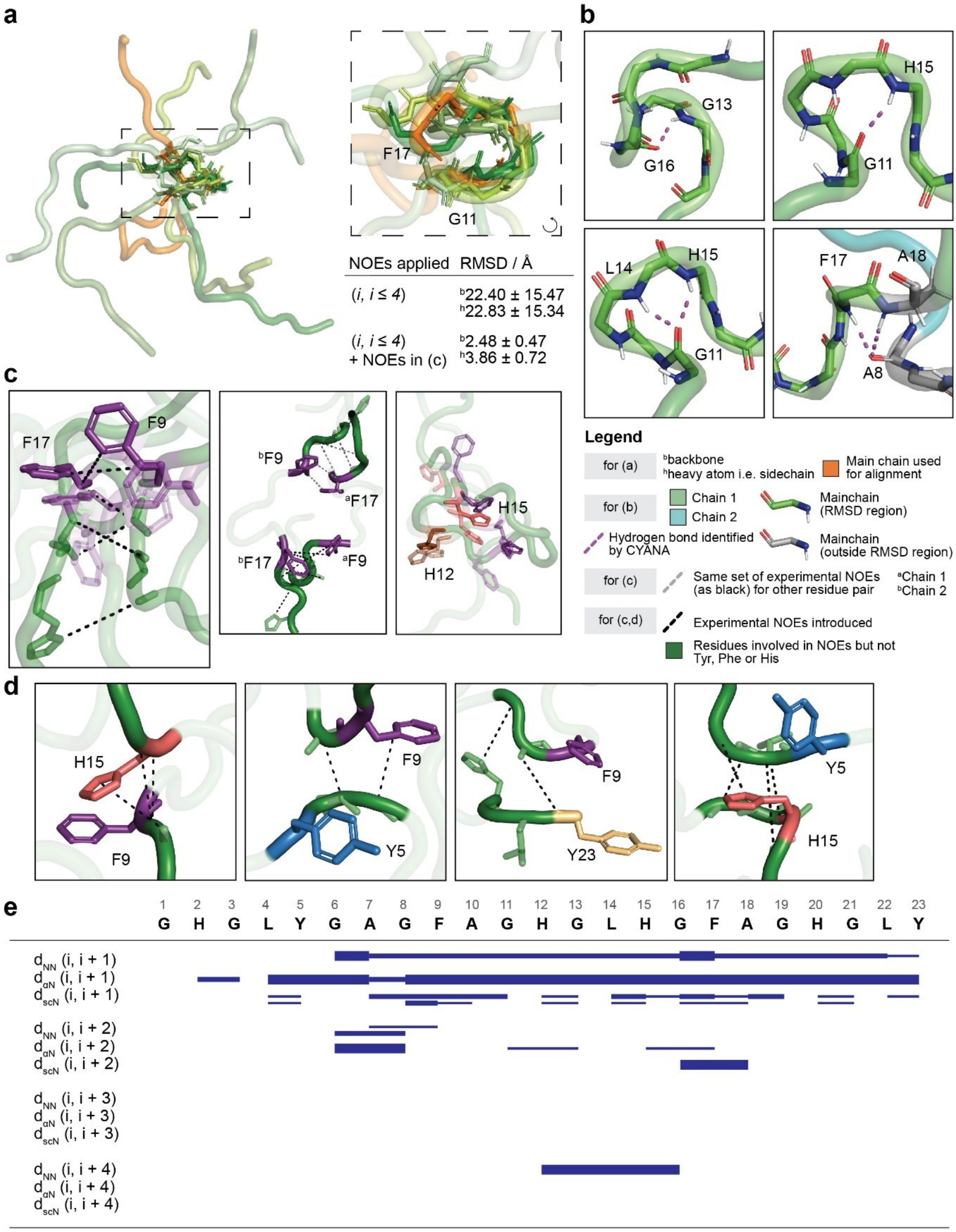

**Supplementary Figure 13. Nonrandom structure of GY23 at pH 3.5. (a)** Overlay of 5 structural conformers from the ensemble calculated. The region used for RMSD calculation and structural alignment is represented using stick representation for the main chain. Top view of the RMSD region is as depicted, with the starting and ending residue labelled and indicated with the direction of the peptide chain. NOEs values reported are direct readouts from CYANA. **(b)** Depiction of hydrogen bonding between mainchain atoms (NH…O) found in several conformers. **(c)** Depiction of F17-F9 interaction pair with experimental NOEs as black dashed lines. Different structural conformers are shown with reduced opacity to highlight the dynamicity of such interactions. An overview of two such pairs is also shown, with the duplicated set of NOEs in grey. Depiction of the spatial arrangement of histidine residues (H12; brown and H15; pink) and the Phe residues **(d)** Depiction of other interactions found from the calculated structures. Structural conformers are not shown for all interactions due to less clarity but are observed to be dynamic. **(e)** Mapping of sequential and short range (up to i, i+4) NOEs at pH 3.5 along peptide chain. Each NOE is indicated by a horizontal bar connecting the proton positions, with the bar thickness proportional to the NOE intensity. $d_{scN}$ indicates the connection between amide protons to any sidechain protons.

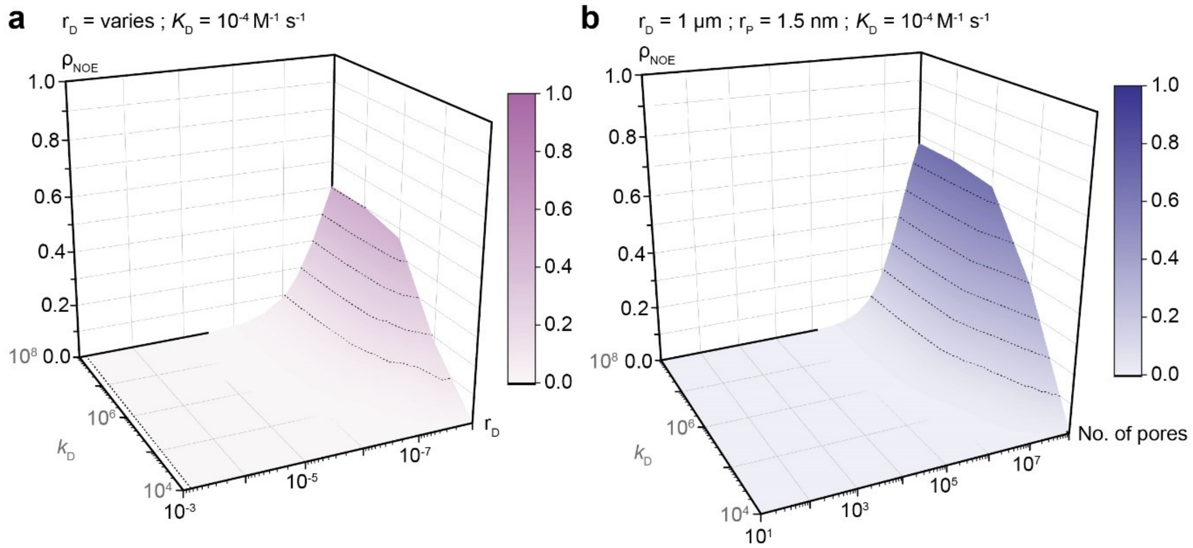


**Supplementary Figure 14. Calculated NOE intensities as function of varying diffusion-limited rate constant, $k_D$ with a fixed dissociation constant, $K_D$ = $10^{-4}$ $M^{-1}$ $s^{-1}$. (a)** NOE intensities are calculated as a function of $k_D$ and droplet radius, $r_D$. **(b)** NOE intensities are calculated as a function of $k_D$ and the number of pores. The radius of the droplet, $r_D$, is fixed as 1 μm, and the pore radius, $r_P$, is predefined at 1.5 nm. Dashed lines on graph plots correspond to an interval of 0.1 on the colour scale as a guide for the eyes. NOE intensity, $\rho_{NOE}$, is defined as the ratio of the intensity of the cross peak to the intensity of the corresponding diagonal peak. Detailed calculations can be found in Section S3.

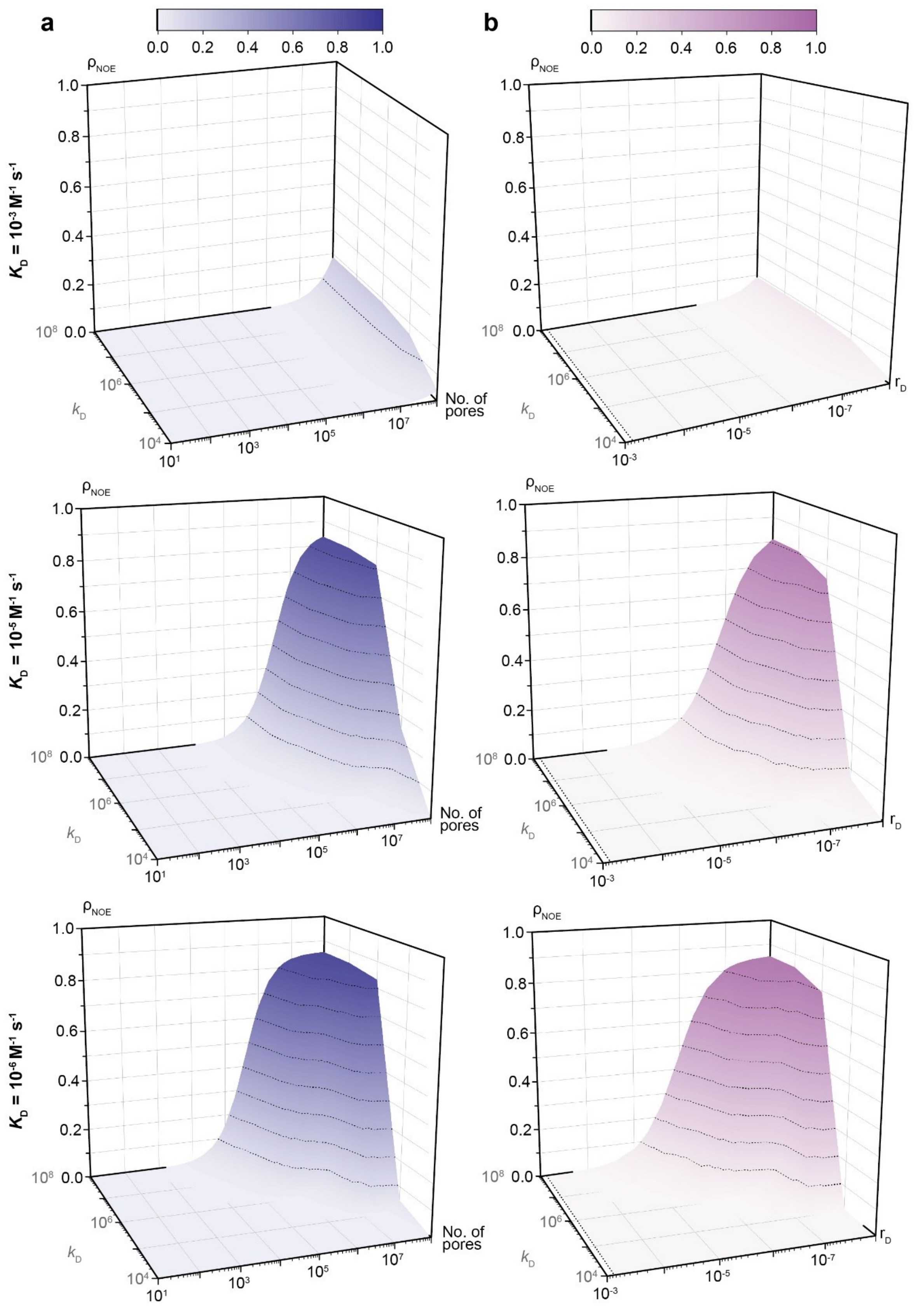


**Supplementary Figure 15. Calculated NOE intensities for other fixed $K_D$ values.** NOE intensities are calculated as a function of the diffusion-limited rate constant, $k_D$ and **(a)** the number of pores or the **(b)** droplet radius. The rendering of graphs is as per Fig. S14.

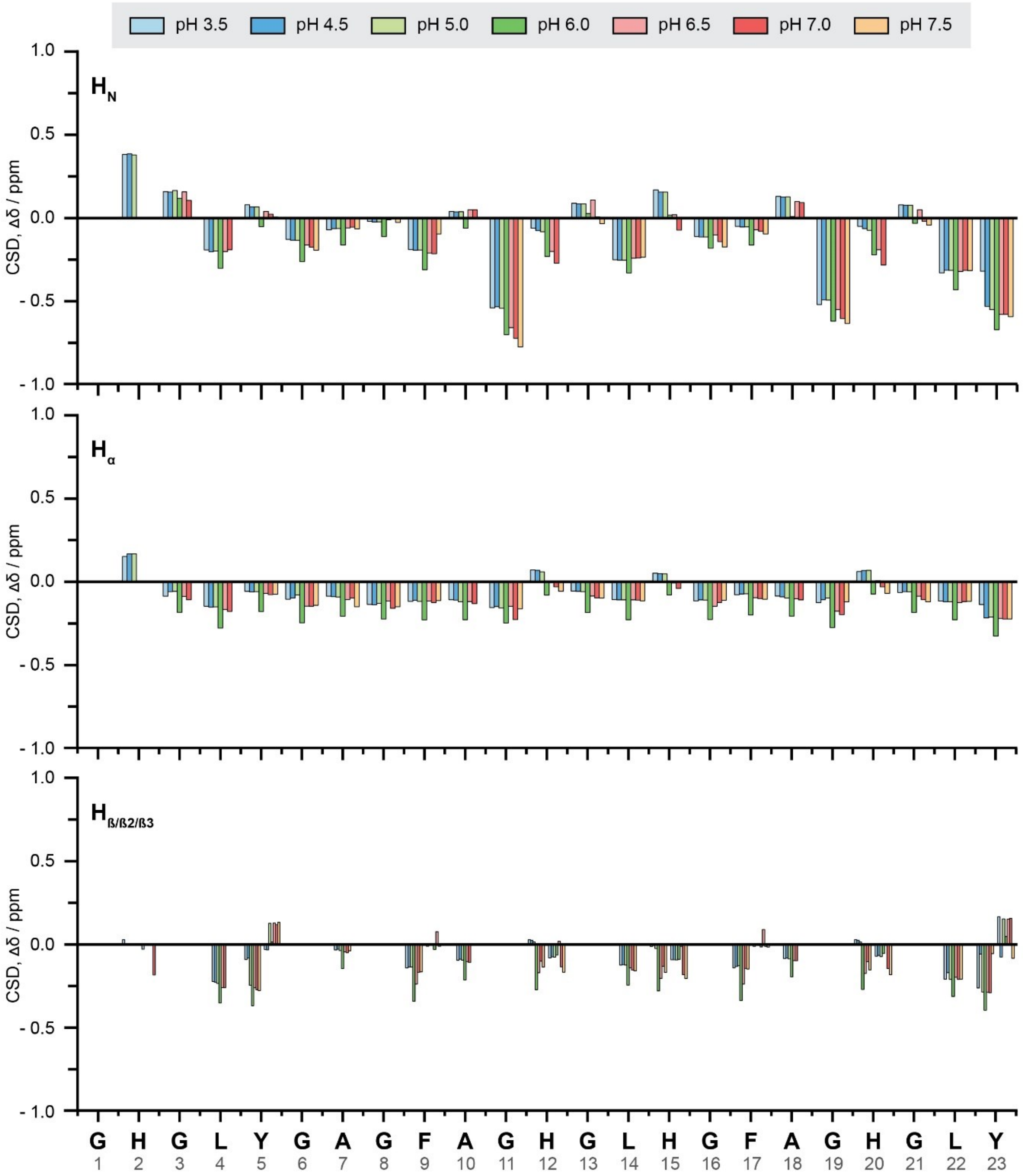


**Supplementary Figure 16. Chemical shift deviations of protons across pH.** Chemical shift deviation values from random coil values[2] obtained for protons ($H_N$, $H_\alpha$, $H_\beta$) across the peptide length. Bar plots are coloured according to the corresponding pH condition.

**Table S1. Derived values from the fitting of power law (coacervate surface) in low Q region of plot in Fig. S11.**

| Time elapsed / min | Value |
|---|---|
| Initial | 3.19 ± 0.05 |
| 5 | 3.26 ± 0.06 |
| 30 | 3.24 ± 0.05 |
| 180 | 3.19 ± 0.22 |
| 240 | 3.19 ± 0.27 |
| 300 | 2.98 ± 0.32 |

**Table S2. List of LogP and surface hydrophobicity values of cargo molecules used in Fig. 6.**

| **Molecule** | **LogP[1]** |
|---|---|
| Carboxyfluorescein | 2.54 |
| Rhodamine B | 1.66 |
| Sulforhodamine B | -0.22 |
| DroProbe | 4.89 |
| Dextran | -1.53 |
| **Protein** | **Surface Hydrophobicity[2]** |
| Insulin | 0.590 |
| Ubiquitin | 0.539 |
| eGFP | 0.566 |
| mCherry | 0.532 |
| Wheat germ agglutin | 0.565 |
| Transferrin | 0.534 |
| Immunoglobulin C | 0.574 |

[1]Small molecules are prepared using ACD ChemSketch Software[3] and uploaded onto ALOGPS 2.1[4-13] online applet for LogP prediction.
[2]Surface hydrophobicity computed using PDBParam[14] and PDB files of proteins, correspondingly depicted in Figure 6.

**Table S3. Calculations for the 3 cases presented in Fig. 7.**

| Case 1 | Case 2 |
| --- | --- |
| Assuming that 50% of peptides are in the dense phase *vs.* the dilute phase and 100% volume of the droplet is occupied by peptides with no overlap,<br><br>Radius of droplet = 1 x $10^{-6}$ m<br>Volume of a droplet = 4.2 x $10^{-18}$ $m^3$<br>Volume of a peptide = 3.0 x $10^{-28}$ $m^3$<br>No. of peptides in a droplet = 1.4 x $10^{10}$<br>No. of peptides in dilute phase = $10^{10}$<br><br>No TrNOESY will occur as there are no exchangeable surfaces. | Imagine a monomolecular layer of peptides at the interface of the droplets which is exchanging with the dilute phase,<br><br>Surface area of droplet = 6.0 x $10^{-19}$ $m^3$<br>No. of peptides at the surface = 2.1 x $10^7$<br>Concentration of available binding sites (as per Section S3)<br>= $(10^7/10^{10})$ x 0.5 x $10^{-3}$<br>= $10^{-6}$ M<br><br>Different orientations of the peptides, i.e., parallel to the surface of the droplet or perpendicular to the surface, as illustrated in Figure 7, were found to give the same magnitude, $10^7$.<br><br>Therefore, the ratio of the concentration of free peptides (ligands) to the available binding sites is $10^{-3}$ to $10^{-6}$, which is 0.1 %. |

| Case 3 |
| --- |
| Imagine a porous structure where each pore is effectively surrounded by an exchangeable monolayer. The surface area to volume ratio thereby increases which will lead to an increase in TrNOE intensity, resulting in observable signals. For example, for a 100x increase in surface area,<br><br>Surface area available = 1.256 x $10^{-9}$<br>No. of peptides at the surface = 2.1 x $10^9$<br>Concentration of available binding sites = $10^{-4}$ M<br><br>The concentration ratio is now $10^{-1}$, which works out to 10%, which is our presumed detectability threshold. |

**Table S4. Buffers used for different pH conditions.**

| pH range | Buffer (50 mM) |
|---|---|
| 4.5, 5.0 | Sodium acetate (acetic acid) |
| 6.0 | Citric acid |
| 7.0 | Sodium phosphate |
| 7.5 | Tris – HCL |

**Table S5. Calculated diffusion constants[1] from DOSY NMR and DECRA Analysis**

| pH | GY23 | $(GY23)_2$ | GW26 |
|---|---|---|---|
| 3.5 | $2.09 \times 10^{-10}$ | $1.48 \times 10^{-10}$ | $2.25 \times 10^{-10}$ |
| 5.8 | $2.20 \times 10^{-10}$ | $1.65 \times 10^{-10}$ | $2.08 \times 10^{-10}$ |
| 6.2 | $2.24 \times 10^{-10}$ | $1.63 \times 10^{-10}$ | $2.07 \times 10^{-10}$ |
| 6.6 | $2.19 \times 10^{-10}$ | $6.54 \times 10^{-10}$ | $2.14 \times 10^{-10}$ |
| 7.0 | $1.95 \times 10^{-10}$,<br>$1.51 \times 10^{-10}$ | $2.11 \times 10^{-10}$ | $2.16 \times 10^{-10}$ |
| 7.4 | $2.22 \times 10^{-10}$ | $5.35 \times 10^{-11}$ | $2.22 \times 10^{-10}$ |
| 7.8 | $2.11 \times 10^{-10}$ | $2.13 \times 10^{-10}$ | $2.35 \times 10^{-10}$ |
| 8.2 | $2.26 \times 10^{-10}$ | $1.90 \times 10^{-10}$ | $2.19 \times 10^{-10}$ |

[1]Units are in $m^2/s$.

**Table S6. Calculated diffusion constant[1] of ubiquitin from DOSY NMR and DECRA Analysis**

| Ubiquitin |
|---|
| $1.60 \times 10^{-10}$ |

[1]Units are in $m^2/s$.

### S1. Experimental details of sample preparation for Small Neutron Angle Scattering (SANS) studies and data processing

#### S1.1 GY23 deuteration and purification for SANS

Deuterated GY23 (dGY23) fusion protein was over-expressed in *E. coli* strain BL21 (DE3) adapted to growth in deuterated minimal medium[15]. The construct of the fusion protein, used as per an earlier study[16], consists of an N terminal HBP-1 protein (original longer sequence of GY23; 112 residues) and a trypsin cleavage site, i.e. Lys residue and GY23 (23 residues). A 1.8 L (final volume) deuterated high cell-density fed-batch fermenter culture was carried out at 30 °C. Feeding with glycerol was started at an $OD_{600}$ value of about 4.35. Expression was induced at an $OD_{600}$ of about 13.4 by addition of IPTG (1 mM final concentration). Cells were harvested at an $OD_{600}$ of 16 yielding 60 g wet weight of perdeuterated cell paste.

Purification was done as previously described[16]. Briefly, the cell paste was resuspended in a lysis buffer (50 mM Tris-HCL, pH 8), supplemented with Protease Inhibitor Cocktail Set I, Calbiochem and passed through a microfluidizer (18000 psi, 4-5 passes, 4°C). The lysate was mixed with ice-cold 100 % acetic acid in a 1:20 volume ratio (acid:lysate) and centrifuged (38000 *g*, 40 min, 4°C). The supernatant was subsequently purified through reverse-phase High Performance Liquid Chromatography (HPLC) with the same purification protocol of the synthetic peptides (see Methods). Desired fractions were identified through mass spectrometry, pooled and freeze-dried to remove traces of acetonitrile. Trypsin cleavage was carried out to cleave GY23 from the fusion protein. Trypsin from the bovine pancreas (TPCK Treated, Sigma Aldrich, Singapore) was dissolved in 50 mM acetic acid to a final concentration of 1 mg/mL. Cleavage was performed at pH 8, 37°C, overnight with shaking at a 1:50 mass ratio (trypsin:peptide). The resultant mixture was again purified through reverse-phase HPLC using the same protocol. Fractions containing dGY23 were identified, pooled and freeze-dried.

#### S1.2 Sample Preparation for SANS and Rationale

Samples were prepared immediately before measurement. For the dilute phase, solutions of 5 mg/ml deuterated (dGY23) in 10 mM acetic acid solution in 42% $D_2O$ were prepared and measured. For the dense phase, the full contrast data GY23 in 100 mM $NaPO_4$ buffer in 100% $D_2O$ was prepared. The goal, though, was to study the hierarchical building blocks present within the coacervate. That was achieved by spiking 15 mg/ml hydrogenous GY23 (hGY23) with 5 mg/ml dGY23 in a 100 mM $NaPO_4$ buffer in 42% $D_2O$. The 42% $D_2O$ was selected to solvent match with the hGY23 SANS signal, allowing the structures of the building blocks within the coacervates to be studied as if in dilute conditions (Fig. S12). The solvent matching efficiency was verified by measuring a sample of 15 mg/ml GY23 in a 100 mM $NaPO_4$ buffer 42% $D_2O$, where no scattering was present within the Q region studied.

In the dilute regime (i.e. under solvent matching), SANS with selective deuteration is a low-resolution method to determine the form factor of the molecule or a group of molecules, which is related to the shape of such particles. As the solution becomes more concentrated, a structure factor may arise that corresponds to the intermolecular interactions between the particles and may give rise to correlation peaks between them due to clustering[17-19]. Using selective deuteration and contrast matching, taking advantage of the large scattering ability between hydrogen and deuterium, the scattering arising from a part of a molecule or a molecule within a complex can be isolated, and this method provided the first structure of protein-nucleic acid complexes in nucleosomes and many protein-protein complexes[20].

**S1.3 Data Processing for SANS**

Data was processed using IgorPro8, on which background subtraction was done (IrenaSAXS macros). Model-free data analysis was performed using the ATSAS suite of programs[21]. The Guinier wizard was used to determine the radius of gyration ($R_g$) by applying a fit in the region to $\log(I(q)) = \log(I(0)) - Q^2 * R_g^2/3$ for $Q * R_g < 0.8$ for extended particles, where I(0) is the forward scattering intensity. The distance distribution module was used to create a pair distribution plot (p(r) vs r) through an indirect Fourier transformation of the raw scattering data since smoothing did not change the shape of the pair distribution function.

The Porod invariant was used to calculate the hydrated volume (V) of the particle following the equation:

$$V = \frac{2\pi^2 I_{exp}^2(0)}{\int_0^{\infty} I(q)q^2 dq}$$

The full contrast data was fitted to a power law using $I(Q) = A * Q^{-n}$ where n is the fractal degrees of freedom and n -= 6- D, where D is the fractal dimension. When D lies between 3 and 4, the scattering can be assigned to a surface mass fractal, which exists only at the boundary of the fractal[22,23].

### S2. Small oligomers are detected at non-coacervating conditions

For dGY23 at pH 3.5, the form factor of the peptide was clearly observed (Fig. 4a) since a clear Guinier region could be identified (a linear relationship on the log $I$ *vs.* $Q^2$ plot, for a qRg < 0.8), suggesting the formation of a defined particle absent from interparticle interactions. From this Guinier region, the $R_g$ was calculated to be roughly 2.5 nm (Table 1). This value was too large to correspond to a single random coil peptide in solution ($R_g = 13$ Å based on Flory's theory[24]), suggesting oligomer formation in acidic conditions. A similar signature to that of the dense phase was observed for the dilute phase in Kratky's plot (Fig. 4b). The Porod volume was calculated, yielding a value consistent with an oligomer composed of 4-5 peptides. Finally, the pair distribution plot calculated from the scattering curve gave a multi-bell-shaped function (Fig. 4c). Such a shape suggested several nodes or sub-domains, perhaps interconnected by disordered regions[25]. However, it was unlikely that the scattering curve arose from a unique monodisperse oligomeric structure but rather from a conformational assembly of different oligomeric structures. In that case, the $p(r)$ *vs.* $r$ function represented the volume fraction-weighted contribution from each population in the ensemble of oligomer structural states, where the presence of multidomain existed. Taken together, the model-free SANS data analysis of dGY23 suggested that oligomeric structures were formed under acidic conditions that contained both globular and extended-like/disordered domains.

Analysis of NOEs from NMR at pH 3.5 indicated the presence of a nonrandom structure of GY23. Structural conformers calculated using CYANA showed that the peptide ends remained conformationally flexible. In contrast, most of the structural features were located in the middle section of the peptide chain, i.e. G11 to F17. While the calculated conformers showed helical elements (mostly single 3-turns) along the peptide chain (Fig. S13b), the absence of *d(i, i+3)* NOEs suggested that these helical structures were not canonical (Fig. S13e). The $H^{\alpha}$ and $H^{N}$ chemical shift deviations (CSD) from random coil values also did not highlight any regions of residues with significant deviations, except for certain residues interspersed throughout the peptide length (Fig. S16). Instead, the sparse *d(i, i+2)* NOEs contrasted against the homogenous distribution of sequential NOEs indicate that the majority of the residues were involved in forming β-structure, specifically β-bridges, comprising of hydrogen bonding between both intra- and interpeptide backbone atoms (Fig. S13b). Remarkably, both long-range, $d(i, i \geq 10)$ and medium-range, $d(i, 4 < i < 10)$ NOEs were also present. These NOEs were carefully studied before being grouped into interaction hubs.

At pH 3.5, interactions revolved around the two Phe residues, F9 and F17 (Fig. S13c), which led to a clear stabilization of the conformers calculated, demonstrated by the significantly reduced root mean square deviation (RMSD) (Fig. S13a). A few NOEs were found between Y5 and F9 as well as Y23 and F9 (Fig. S13d), indicating that these interactions might be weak. Several more NOEs were found between F9 and H15, and surprisingly, H15 and Y5, though the latter may be due to coincidental proximity as the sidechain of H15 faced the same side as the Phe residues in most conformers calculated (Fig. S13c), possibly explaining the observed 'preference' for H15 over other histidines. Clustering of the aromatic rings[26], as well as interactions between protonated histidine and aromatic rings[27], have been shown to lead to the stabilization of proteins and peptides at low pH. This was also evident in the energy values computed for the interacting pairwise configurations in Fig. S8. The calculated diffusion constant (Table S5) for GY23 from DOSY NMR experiments also supported the presence of small oligomers (around 3 peptides on average) at low pH, as the derived value was similar to that obtained for ubiquitin (Table S6).

Interestingly, although $(GY23)_2$ had a similar distribution of interproton resonances at pH 3.5 (Fig. S10b), significantly fewer resonances were found for GW26, suggesting a more extended, water-exposed conformation of the peptide, possibly facilitating its phase separation at low pH, albeit under high salt conditions. The DOSY diffusion constants (Table S5) calculated for these two peptides were also similar to that for GY23, though slightly reduced for $(GY23)_2$, indicating that these peptides also formed small-sized oligomers at low pH.

### S3. Formulation using analytical expressions for calculation of TrNOE intensities

#### S3.1 Transferred NOEs in the coacervating peptide systems.

The concentration of binding sites available for free peptides in solution in chemical exchange with the peptides locked in the droplets is estimated by assuming (i) a monomolecular surface of the droplet mediating the exchange, (ii) the geometric shape of the exchanging peptide can be approximated by a box with sides $d_{xyz}$ = 0.5, 1.2 and 0.5 nm representing two lateral and one depth dimensions. The number of the peptide binding sites on the monomolecular surface of a droplet, $N_B$, of the radius $r_D$ and the number of the peptides forming the interior of the dense droplet, $N_V$, are given by Eqs. 1A and B.

$$N_B = \frac{4\pi(r_D)^2}{d_x d_y} \tag{1A}$$

$$N_V = \frac{\frac{3}{4}\pi(r_D)^3}{d_x d_y d_z} \tag{1B}$$

Several studies[28,29] in literature have estimated that the number of peptides in the interior of the droplet exceeds the number of the surface layer forming peptides by the order of $10^2$ to $10^4$, fitting our calculations though we note that this magnitude can possibly vary due to droplet size. The total concentration of the peptide binding sites is given by Eq. 2:

$$[B]_T = \frac{N_B}{N_V}[1 - c_{free}][P]_T \tag{2}$$

where $c_{free}$ is the peptide partition coefficient between the solvent and droplets and $[P]_T$ is the total peptide concentration. These parameters are experimentally accessible using 1D $^1$H NMR spectra and are reported in Fig. 1.

In the case that each individual binding event of the peptide to its surface binding site does not cause significant allosteric restructuring in the droplet (and thus, all binding events are essentially independent), the exchange process of the peptides between solution and droplets can be described as a bimolecular reaction given by Eq. 3

$$P + B \leftrightarrow PB \tag{3}$$

with the associated equilibrium constant $K_D$ , the kinetic rates $k_{on}$ and $k_{off}$ . The equilibrium concentrations [P], [B] and [PB] are available as a function of the kinetic rates and total peptide concentration[30]. For bimolecular reactions, the reaction rate depends on the number of collisions per unit time, which is proportional to the product of the concentrations of the reactants. For the reaction in Eq. 3, the general rate law for the association and dissociation molecular fluxes[31] is as follows

$$J_1 = \kappa[P]([B] + [PB]) \tag{4A}$$

$$J_{-1} = k_{off}[PB] \tag{4B}$$

where κ is the reaction rate coefficient. Since the molecular collisions of the peptide with the unoccupied binding site B as well as with the "occupied" binding site PB result in the peptide association, the [B] is extended with [PB] in Eq. 4A.

The elements of the corresponding kinetic matrix $\boldsymbol{K}$ for the populations of the pair of proton spins (A-X) residing in the peptide exchanging between solvent and droplet, respectively, are given by Eqs. 5:

$$k_1 = \kappa([B] + [PB]) \tag{5A}$$

$$k_{-1} = k_{off} \tag{5B}$$

Assuming that the association reaction is primarily diffusion-controlled (or diffusion-limited) in which the reaction rate is equal to the rate of transport of the reactants through the solution and that the intrinsic association rate (formation of productive intermolecular interactions) is much faster than the diffusion rate and the intermolecular forces are weak, κ can be approximated by the corresponding diffusion-limited rate constant $k_D$. Although $k_D$ can be recalculated from the translation diffusion rates as measured by $^1$H NMR DOSY experiments using Smoluchowski's theory[32,33], the realistic $k_D$ estimation may require knowledge of the binding funnel, distribution of reactive sites and required steric specificities between associating peptide and its binding sites[32,33]. In the current analysis, we opted to explore the realistic range typically exhibited by the proteins of similar size (comparable to small peptide oligomers), e.g. $k_D$ in the range of $10^5$–$10^6$ $M^{-1}$ $sec^{-1}$. A wider range from $10^4$ to $10^8$ $M^{-1}$ $sec^{-1}$ was still explored to cover other possible oligomeric species.

Analytical expressions for the Transferred NOESY cross-peak intensities of a two-spin system (A-X) in the absence of ligand-protein cross relaxation[34,35] were used to estimate the ratio of the NOESY cross-peak intensity to the corresponding diagonal peak intensity for the exchanging peptide between solvent and the droplet via binding to its surface. These auto relaxation-normalized TrNOEs were explored numerically as a function of the droplet radius, as shown in (Fig. 5b) and Fig. S14 and S15 (or the number of spherical pores with the predefined radius $r_P$ within a droplet with an overall predefined radius), the range of diffusion-controlled reaction rates $k_D$ and the equilibrium dissociation rate constant $K_D$ estimated as a concentration of the peptide at selected LLPS conditions when ½ of the total peptides in solution were partitioned into the droplet phase and reported as the LLPS phase diagram boundaries in Fig. 1. To cover the different phase separation propensities across the peptides, a range was explored. The MATLAB code used in the simulation is listed below.

#### S3.2 MATLAB Code

Loops were used to calculate the data for the respective graph plots. Due to the numerous terms used, only the base codes are shown here, which can be readily adapted to varying cases, such as predefining the droplet radius and pore radius for the calculation of TrNOE intensities as a function of the number of pores (code variation not shown here). The exact code used to cover the different cases explored in this work can be provided upon request. *G* and *H* here are equivalent to *P* and *B,* respectively.

```
%% Setting of general parameters
```

```
Pi = 3.14159
tmix =0.2; % s, NOESY mixing time
tcW = 10*10^-9; % free ligand
wW = 700.0*10^6; % Proton Larmor frequency
r = 3.0*10^-10; % distance between two 1H in the ligand
Kd = 10^-4 ; % M, dissociation constant

%% Finding the number of binding sites on the surface of dense droplet as
function of its radius, Rdroplet
dx = 5*10^-10; % lateral size x of the gy23 as ligand - 5A
dy = 12*10^-10; % lateral size y of the gy23 as ligand - 12A
dz = 5*10^-10; % depth size z of the gy23 as ligand - 5A
Cpartition = 0.5; % partition coeff of gy23 between liquid and dense phases, %
of free ligand left at coacervation
Rdroplet = 1*10^-6;  % meters, radius of a droplet

Ligand_concentration_at_noLLPS = 1*10^-3; % 1 mM of all ligand at no
coacervation
Ltotal = Ligand_concentration_at_noLLPS*Cpartition % 0.5 mM of total free
ligand at coacervation
%i.e. not partitioned peptides

Sdroplet = 4*Pi*(Rdroplet)^2;  % % surface area of a droplet
Nb = Sdroplet/(dx*dy); % number of binding sites on the surface of a droplet
Vdroplet = 4/3*Pi*(Rdroplet)^3;  % % surface area of a droplet
Nv = Vdroplet/(dx*dy*dz); %no. of peptides making up the droplet
Ptotal = Nb/Nv*Ligand_concentration_at_noLLPS*(1-Cpartition) % concentration
of binding sites
NfNb = Ltotal/Ptotal % ratio of free ligands to number of binding sites

%% Kinetics of exchange to calculate k, and kprime.
Kon_diffusion_limited = 10^6; % M-1 s-1.
Khg = 1/Kd; % Units M^-1
%% Khg = [HG]/[[H][G]] and Kd is [[reactant][reactant]]/[product]
Ht = Ptotal; % host total - concentration of binding sites - units M
Gt = Ltotal; % guest total - free ligands - units M

H = (Khg*(Ht-Gt)-1 +  sqrt( (Khg*(Ht-Gt)-1)^2+4*Khg*Ht )   )/(2*Khg)
HG =(Khg*(Ht+Gt)+1 -  sqrt( (Khg*(Ht-Gt)-1)^2+4*Khg*Ht )   )/(2*Khg)
G = (Khg*(Gt-Ht)-1 +  sqrt( (Khg*(Ht-Gt)-1)^2+4*Khg*Ht )   )/(2*Khg)

Koff = Kd*Kon_diffusion_limited % units = [M] * [M-1][s-1]
Kexchange = KonL + Koff;
Percent_L_detectable_by_nmr = G/Gt
HGpopulation = HG/(H+G)
k = Kon_diffusion_limited*(H+HG)
k_prime = Koff % units = [s-1]

%% Calculations of NOE Intensities
[rho,sigma] = cross_relaxation_rates(3*10^-9, 700.0*10^6, 3.0*10^-10); % (tcW,
wW, r) for free ligand
[rho_prime,sigma_prime] = cross_relaxation_rates(300*10^-9, 700.0*10^6,
3.0*10^-10); % (tcW, wW, r) for bound ligand
s = rho_prime +k_prime - sigma_prime;
r = rho +k - sigma;
p = rho+k+sigma;
```

```
q = rho_prime+k_prime+sigma_prime;
F = r+s;
E = r-s;
D = sqrt(E^2+4*k*k_prime);
C = p+q;
B = p-q;
A = sqrt(B^2+4*k*k_prime);
lambda1 = (C-A)/2;
lambda2 = (F-D)/2;
lambda3 = (C+A)/2;
lambda4 = (F+D)/2;

I_AA = (1-B/A)*exp(-lambda1*tmix)+(1-E/D)*exp(-lambda2*tmix)+(1+B/A)*exp(-
lambda3*tmix)+(1+E/D)*exp(-lambda4*tmix);
I_AX = (1-B/A)*exp(-lambda1*tmix)-(1-E/D)*exp(-lambda2*tmix)+(1+B/A)*exp(-
lambda3*tmix)-(1+E/D)*exp(-lambda4*tmix);

I_AA
I_AX
SV = Sdroplet/Vdroplet

function [rho,sigma] = cross_relaxation_rates(tcW, wW, r)
distance_between_protons_in_water = 0.96*10^-10; % A

K = 1.02*10^10; % s^-2, for relaxation of two protons in water
W1 = 3*K*tcW/(1+tcW^2*wW^2)*(distance_between_protons_in_water)^6 / (r^6);
W0 = 2*K*tcW*(distance_between_protons_in_water)^6 / (r^6);
W2 = 12*K*tcW/(1+4*tcW^2*wW^2)*(distance_between_protons_in_water)^6 / (r^6);
rho = 2*W1+W0+W2; % checked
sigma = W2-W0;
end
```
